# Deep mutational scanning of SARS-CoV-2 receptor binding domain reveals constraints on folding and ACE2 binding

**DOI:** 10.1101/2020.06.17.157982

**Authors:** Tyler N. Starr, Allison J. Greaney, Sarah K. Hilton, Katharine H.D. Crawford, Mary Jane Navarro, John E. Bowen, M. Alejandra Tortorici, Alexandra C. Walls, David Veesler, Jesse D. Bloom

## Abstract

The receptor binding domain (RBD) of the SARS-CoV-2 spike glycoprotein mediates viral attachment to ACE2 receptor, and is a major determinant of host range and a dominant target of neutralizing antibodies. Here we experimentally measure how all amino-acid mutations to the RBD affect expression of folded protein and its affinity for ACE2. Most mutations are deleterious for RBD expression and ACE2 binding, and we identify constrained regions on the RBD’s surface that may be desirable targets for vaccines and antibody-based therapeutics. But a substantial number of mutations are well tolerated or even enhance ACE2 binding, including at ACE2 interface residues that vary across SARS-related coronaviruses. However, we find no evidence that these ACE2-affinity enhancing mutations have been selected in current SARS-CoV-2 pandemic isolates. We present an interactive visualization and open analysis pipeline to facilitate use of our dataset for vaccine design and functional annotation of mutations observed during viral surveillance.

## Introduction

The SARS-related (sarbecovirus) subgenus of betacoronaviruses comprises a diverse lineage of viruses that circulate in bat reservoirs and spill over into other mammalian species (Bolles et al., 2011; Cui et al., 2019). Sarbecoviruses initiate infection by binding to receptors on host cells via the viral spike surface glycoprotein. The entry receptor for SARS-CoV-1 and SARS-CoV-2 is the human cell-surface protein angiotensin converting enzyme 2 (ACE2), and the receptor binding domain (RBD) of spike from both these viruses binds ACE2 with high affinity (Hoffmann et al., 2020; Letko et al., 2020; Li et al., 2003; Walls et al., 2020; Wrapp et al., 2020a). Because of its key role in viral entry, the RBD is a major determinant of cross-species transmission and evolution (Becker et al., 2008; Frieman et al., 2012; Letko et al., 2020; Li, 2008; Li et al., 2005b; Qu et al., 2005; Ren et al., 2008; Sheahan et al., 2008a, 2008b; Wu et al., 2012). In addition, the RBD is the target of the most potent anti-SARS-CoV-2 neutralizing antibodies identified to date (Cao et al., 2020; Ju et al., 2020; Pinto et al., 2020; Rogers et al., 2020; Seydoux et al., 2020; Shi et al., 2020; Wu et al., 2020; Zost et al., 2020), and several promising vaccine candidates consist solely of adjuvanted RBD protein (Chen et al., 2020a, 2020b; Quinlan et al., 2020; Ravichandran et al., 2020; Zang et al., 2020).

Despite its important function, the RBD is one of the most variable regions in sequence alignments of sarbecoviruses (Hu et al., 2017), reflecting the complex selective pressures shaping its evolution (Demogines et al., 2012; Frank et al., 2020; MacLean et al., 2020). Furthermore, RBD mutations have already appeared among SARS-CoV-2 pandemic isolates, including some near the ACE2-binding interface—but their impacts on receptor recognition and other biochemical phenotypes remain largely uncharacterized. Therefore, comprehensive knowledge of how mutations impact the SARS-CoV-2 RBD would aid efforts to understand the evolution of this virus and guide the design of vaccines and other countermeasures.

To address this need, we used a quantitative deep mutational scanning approach (Adams et al., 2016; Fowler and Fields, 2014; Weile and Roth, 2018) to experimentally measure how all possible SARS-CoV-2 RBD amino-acid mutations affect ACE2-binding affinity and protein expression levels (a correlate of protein folding stability). The resulting sequence-phenotype maps illuminate the forces that shape RBD evolution, quantify the constraint on antibody epitopes, and suggest that purifying selection is the main force acting on RBD mutations observed in human SARS-CoV-2 isolates to date. To facilitate use of our measurements in immunogen design and viral surveillance, we provide interactive visualizations, an open analysis pipeline, and complete raw and processed data.

## Results

### Yeast display of RBDs from SARS-CoV-2 and related sarbecoviruses

To enable rapid functional characterization of thousands of RBD variants, we developed a yeast surface-display platform for measuring expression of folded RBD protein and its binding to ACE2 (Adams et al., 2016; Boder and Wittrup, 1997). This platform enables RBD expression on the cell surface of yeast (Figure 1B), where it can be assayed for ligand-binding affinity or protein expression levels, a close correlate of protein folding efficiency and stability (Kowalski et al., 1998a, 1998b; Shusta et al., 1999). Because yeast have protein-folding quality control and glycosylation machinery similar to mammalian cells, they add N-linked glycans at the same RBD sites as human cells (Chen et al., 2014), although these glycans are more mannose-rich than mammalian-derived glycans (Hamilton et al., 2003).

**Figure 1.**
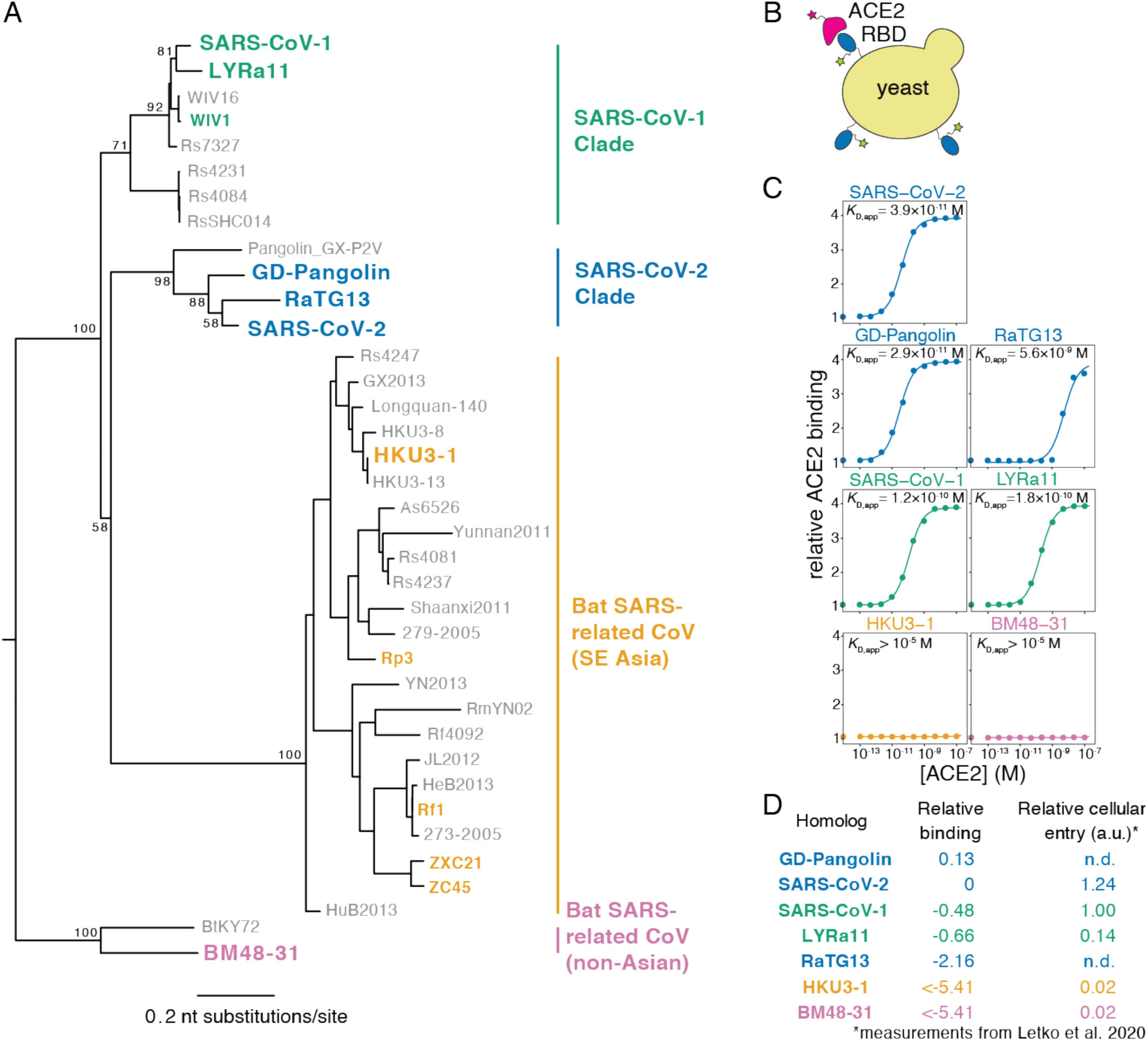
Yeast display of RBDs from SARS-CoV-2 and related sarbecoviruses. (A) Maximum likelihood phylogenetic tree of sarbecovirus RBDs. RBDs included in the present study are in bold text colored by RBD clade. Node labels indicate bootstrap support. (B) RBD yeast surface display enables fluorescent detection of RBD surface expression and ACE2 binding. (C) Yeast displaying the indicated RBD were incubated with varying concentrations of human ACE2, and binding was measured via flow cytometry. Binding constants are reported as *K*_D,app_ from the illustrated titration curve fits. (D) Comparison of yeast display binding with previous measurements of the capacity of viral particles to enter ACE2-expressing cells. Relative binding is Δlog10(*K*_D,app_) measured in the current study; relative cellular entry is infection of ACE2-expressing cells by VSV pseudotyped with spike containing the indicated RBD, reported by Letko et al. (Letko et al., 2020) in arbitrary luciferase units relative to SARS-CoV-1 RBD; n.d. indicates not determined by Letko et al.

To validate the yeast-display platform, we selected RBDs from the Wuhan-Hu-1 SARS-CoV-2 isolate and six related sarbecoviruses (Figure 1A). These other sarbecoviruses include the closest known relatives of SARS-CoV-2 from bats and pangolins (RaTG13 and GD-Pangolin), SARS-CoV-1 (Urbani strain) and a close bat relative (LYRa11), and two more distantly related bat sarbecoviruses (BM48-31 and HKU3-1). Based on prior work, all of these RBDs are expected to bind human ACE2 except those from BM48-31 and HKU3-1 (Lam et al., 2020; Letko et al., 2020; Shang et al., 2020). We cloned the RBDs into a vector for yeast cell surface display, induced RBD expression, and incubated with a fluorescent antibody targeting a C-terminal epitope tag and varying concentrations of fluorescently labeled human ACE2 (Figure 1B). We then used flow cytometry to measure RBD surface expression levels and ACE2 binding across 11 ACE2 concentrations, enabling the calculation of a dissociation constant for the binding of each RBD to ACE2 (Figure 1C). Because we used ACE2 in its native dimeric form (Yan et al., 2020), we refer to the measured constants as apparent dissociation constants (*K*_D,app_) which are affected by binding avidity. We report log binding constants Δlog10(*K*_D,app_) relative to the wildtype SARS-CoV-2 RBD, polarized such that a positive value reflects stronger binding (Figure 1D).

All RBDs expressed well and exhibited ACE2 binding affinities consistent with prior knowledge. We measure *K*_D,app_ = 3.9×10^−11^ M for the SARS-CoV-2 RBD (Figure 1C), which as expected is tighter than affinities reported for monomeric ACE2 (Shang et al., 2020; Walls et al., 2020; Wrapp et al., 2020a) due to avidity effects caused by our use of native dimeric ACE2. Consistent with previous studies (Shang et al., 2020; Walls et al., 2020; Wrapp et al., 2020a), the SARS-CoV-1 RBD binds ACE2 with lower affinity than SARS-CoV-2 (Figures 1C,D). The SARS-CoV-1-related bat strain LYRa11 binds with even lower affinity, while the more distant bat RBDs (HKU3-1 and BM48-31) have no detectable binding. These measurements are consistent with the ability of these RBDs to enable viral particles to enter cells expressing human ACE2 (Letko et al., 2020) (Figure 1D), validating the relevance of our binding measurements to viral infection. Within the newly described SARS-CoV-2 clade, GD-Pangolin binds ACE2 with slightly higher affinity than SARS-CoV-2, while the bat isolate RaTG13 binds with two orders of magnitude lower affinity, consistent with prior qualitative reports (Shang et al., 2020). These results validate our yeast surface display platform for RBD affinity measurements, and map variation in affinity for human ACE2 within the SARS-CoV-2 clade and the broader sarbecovirus subgenus.

### Deep mutational scanning of all amino-acid mutations to the SARS-CoV-2 RBD

We next integrated the yeast-display platform with a deep mutational scanning approach to determine how all amino-acid mutations to the SARS-CoV-2 RBD impact expression and binding affinity for ACE2. We constructed two independent mutant libraries of the RBD using a PCR-based mutagenesis method that introduces all 19 mutant amino acids at each position (Bloom, 2014). To facilitate sequencing and obtain linkage among amino-acid mutations within a single variant, we appended 16-nucleotide barcode sequences downstream of the coding sequence (Hiatt et al., 2010), bottlenecked each library to ∼100,000 barcoded variants, and linked each RBD variant to its barcode via long-read PacBio SMRT sequencing (Matreyek et al., 2018) (Figure S1A). By examining the concordance of RBD variant sequences for barcodes sampled by multiple PacBio reads, we validated that this process correctly determined the sequence of >99.8% of the variants (Figure S1B). RBD variants contained an average of 2.7 amino-acid mutations each, with the number of mutations per variant roughly following a Poisson distribution (Figure S1C). Our libraries covered 3,804 of the 3,819 possible RBD amino-acid mutations, of which 95.7% were present as the sole amino-acid mutation in at least one barcoded variant (Figures S1D,E). To provide internal standards for our measurements, we spiked the mutant libraries with a barcoded panel of 11 unmutated sarbecovirus RBD homologs (strains shown in color in Figure 1A), including those tested in the isogenic assays in Figure 1C.

To determine how mutations affect RBD protein expression and ACE2 binding, we combined fluorescent-activated cell sorting (FACS) with deep sequencing of the variant barcodes (Adams et al., 2016; Peterman and Levine, 2016). To measure expression, we fluorescently labeled RBD protein on the yeast surface via a C-terminal epitope tag and used FACS to collect ∼15 million cells from each library, partitioned into four bins from low to high expression (Figures 2A, S2A). We sequenced the barcodes from each bin and reconstructed each variant’s mean fluorescence intensity (MFI) from its distribution of read counts across sort bins. We represent expression as Δlog(MFI) relative to the unmutated SARS-CoV-2 RBD, such that a positive Δlog(MFI) indicates increased expression. To measure ACE2-binding affinity, we incubated yeast libraries that had been pre-sorted for RBD expression with 16 concentrations of fluorescently labeled ACE2 (10^−6^ to 10^−13^ M, plus no ACE2), and used FACS to collect >5 million RBD+ yeast cells at each ACE2 concentration, partitioned into 4 bins from low to high ACE2 binding (Figures 2B, S2B). We again sequenced the barcodes from each bin, reconstructed the mean ACE2 binding of each variant at each ACE2 concentration, and used the resulting titration curves to infer dissociation constants *K*_D,app_ (Figure S2C), which we represent as Δlog_10_(*K*_D,app_) relative to the unmutated SARS-CoV-2 RBD, with positive values indicating stronger binding.

**Figure 2.**
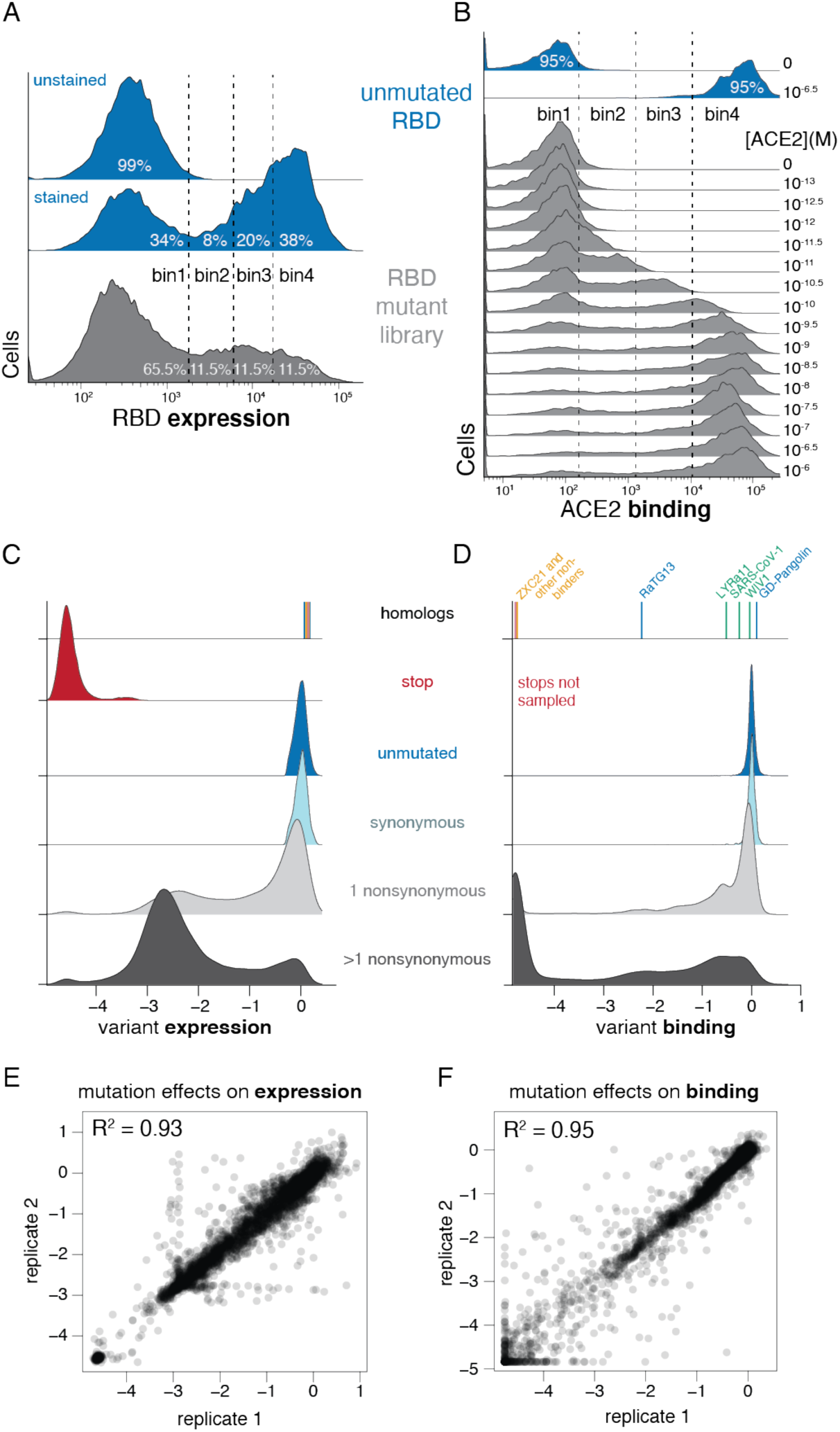
Deep mutational scanning of the SARS-CoV-2 RBD. (A, B) FACS approach for deep mutational scans for expression (A) and binding (B). Cells were sorted into four bins from low to high expression or binding signal, with separate sorts for each ACE2 concentration. The frequency of each library variant in each bin was determined by Illumina sequencing of the barcodes of cells collected in that bin, enabling reconstruction of per-variant expression and binding phenotypes. Bin boundaries were drawn based on distributions of expression or binding for unmutated SARS-CoV-2 controls (blue), and gray shows the distribution of library variants for library replicate 1 in these bins. See also Figure S2. (C, D) Distribution of library variant phenotypes for expression (C) and binding (D), with variants classified by the types of mutations they contain. Internal control RBD homologs are indicated with vertical lines giving the measured expression or binding phenotype, colored by clade as in Figure 1A. Stop-codon-containing variants were purged by an RBD+ pre-sort prior to ACE2 binding deep mutational scanning measurements, and so are not sampled in (D). (E, F) Correlation in single-mutant effects on expression (E) and binding (F), as determined from independent mutant library replicates. See also Figure S3.

These high-throughput measurements of expression and ACE2 binding were consistent with expectations about the effects of mutations. RBD variants containing stop codons universally failed to express folded protein (Figure 2C). Unmutated variants and those with only synonymous mutations had a tight distribution of neutral expression and binding measurements (Figure 2C,D). Variants containing amino-acid mutations had a wide range of expression and binding phenotypes, with variants containing just one mutation tending to have more mild functional defects than those with multiple mutations (Figure 2C,D). These trends are consistent with the well-established fact that most amino-acid mutations are deleterious to protein folding or function (Soskine and Tawfik, 2010)—however, some variants with amino-acid mutations exhibit expression or binding that is comparable or even slightly higher than the parental SARS-CoV-2 RBD. The panel of RBD homologs from other sarbecovirus strains all expressed well but exhibited a wide range of ACE2 binding affinities (Figure 2C,D, Supplemental File 1), as expected from the fact that only some are derived from viruses that can enter cells using human ACE2 (Letko et al., 2020).

These measurements show that the RBD possesses considerable mutational tolerance (Figure 2C, D). For instance, 46% of single amino-acid mutations to SARS-CoV-2 RBD maintain an affinity to ACE2 that is at least as high as that of SARS-CoV-1, suggesting that there is a substantial mutational space consistent with sufficient affinity to maintain human infectivity. Many single amino-acid mutants also maintain expression comparable to that of unmutated SARS-CoV-2, indicating that a large mutational space is compatible with properly folded RBD protein.

We next aggregated the measurements on all variants in our libraries to quantify the effects of individual amino-acid mutations. Because many variants contain multiple mutations, we used global epistasis models to determine the effects of individual mutations from both singly and multiply mutated variants (Otwinowski et al., 2018) (Figure S3). The resulting single-mutant Δlog(MFI) and Δlog_10_(*K*_D,app_) measurements correlated well between the independent library duplicates (R^2^ = 0.93 and 0.95, respectively; Figures 2E,F). Throughout the rest of this paper, we report single mutant effects as the average of the two libraries. Overall, we obtained expression measurements for 99.5% and binding measurements for 99.6% of all 3,819 single amino-acid mutations to the RBD.

### Visualization and validation of sequence-to-phenotype maps

The complete measurements of how amino-acid mutations affect expression and ACE2 binding represent rich sequence-to-phenotype maps for the RBD. These maps are especially informative when interpreted in the context of the RBD’s structure and evolution. To facilitate such interpretation, we visualize the data in several ways. Figure 3 provides heatmaps that show how each mutation affects expression or ACE2 binding, with sites annotated by whether they contact ACE2, their relative solvent accessibility, and their amino-acid identities in SARS-CoV-2 and SARS-CoV-1. Interactive versions of these heatmaps are in Supplemental File 2 and at https://jbloomlab.github.io/SARS-CoV-2-RBD_DMS, and enable zooming, subsetting of positions by functional annotations, and mouse-selection based readouts of numerical measurements for individual mutations. As an alternative representation of the data, Figure S4 provides logo plots that enable side-by-side comparison of how mutations affect expression and ACE2 binding. Finally, interactive structure-based visualizations using the `dms-view` tool (Hilton et al., 2020) are linked at https://jbloomlab.github.io/SARS-CoV-2-RBD_DMS/structures/, and project the effects of mutations onto a crystal structure of the ACE2-bound RBD (Lan et al., 2020) and a cryo-EM structure of the full spike ectodomain (Walls et al., 2020). The underlying raw data in these maps are in Supplemental File 3.

**Figure 3.**
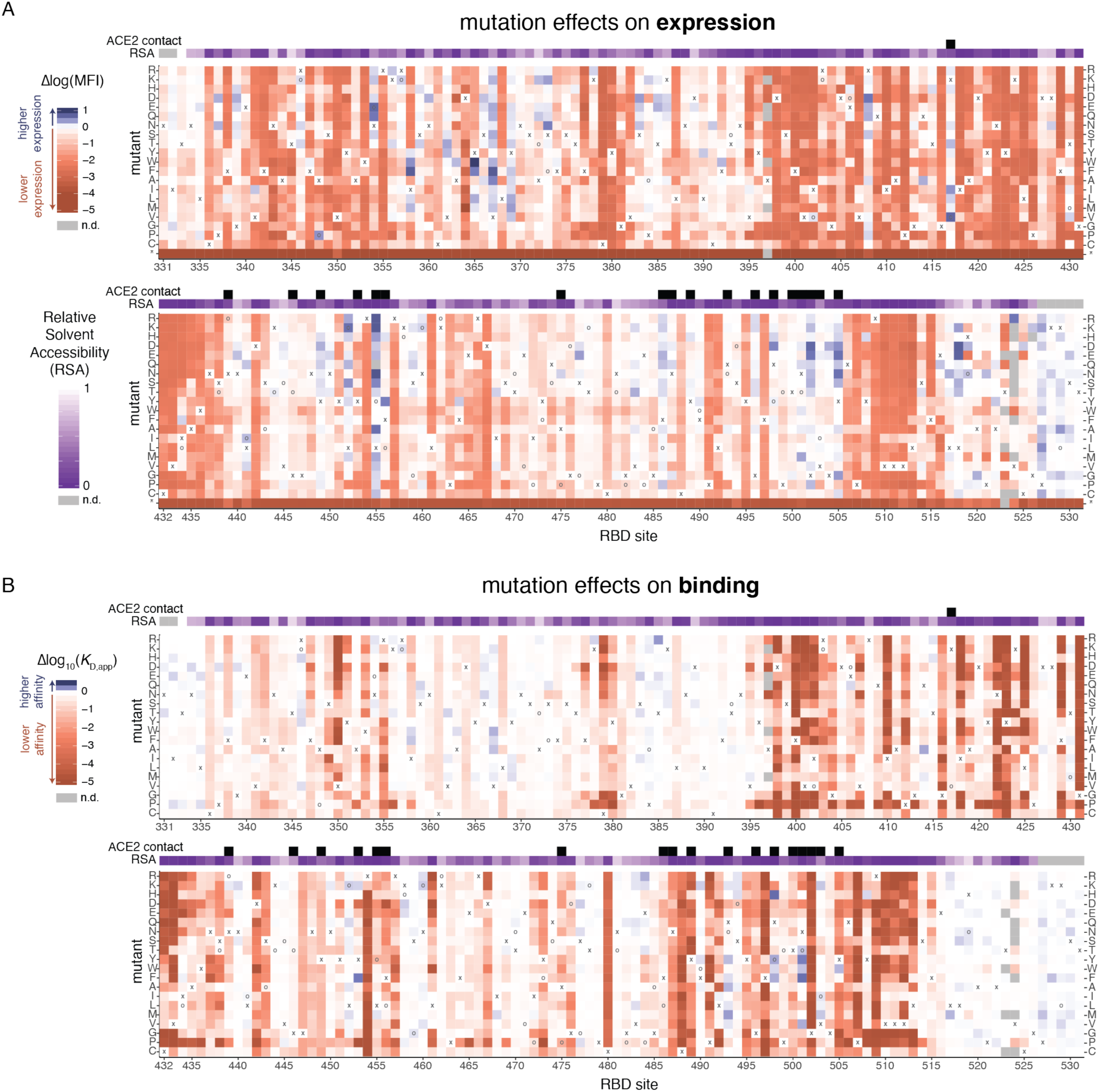
Sequence-to-phenotype maps of the SARS-CoV-2 RBD. (A, B) Heatmaps illustrating how all single mutations affect RBD expression (A) and ACE2 binding affinity (B). Interactive versions of these heatmaps are at https://jbloomlab.github.io/SARS-CoV-2-RBD_DMS and in Supplemental File 2. Squares are colored by mutational effect according to scale bars on the left, with red indicating deleterious mutations. The SARS-CoV-2 amino acid is indicated with an ‘x’, and when the SARS-CoV-1 amino acid is different it is indicated with an ‘o’. The top overlay uses black boxes to indicate residues that contact ACE2 in the SARS-CoV-2 (PDB 6M0J) or SARS-CoV-1 (PDB 2AJF) crystal structures. The purple overlay represents the relative solvent accessibility (RSA) of a residue in the ACE2-bound SARS-CoV-2 crystal structure. See also Figure S4.

The sequence-phenotype maps reveal tremendous heterogeneity in mutational constraint across the RBD. Many sites are highly tolerant of mutations with respect to one or both of expression and ACE2 binding, while other sites are highly constrained to the wildtype amino acid in SARS-CoV-2. A substantial number of sites (e.g., sites 382 to 395) are quite tolerant of mutations with respect to ACE2 binding, but are constrained with respect to expression—consistent with folding and stability being global constraints common to many sites (Fane et al., 1991; Poteete et al., 1997). There are also a handful of sites where ACE2 binding imposes strong constraints but expression does not (e.g. sites 489, 502, and 505). Moreover, at some sites there are mutations that clearly enhance expression or ACE2-binding affinity (blue colors in Figure 3).

To validate key features of the sequence-phenotype maps, we performed experiments to confirm the dynamic range of our assays and their relevance in the context of purified RBD protein and pseudotyped lentiviral particles (Figure 4). We first re-cloned and tested a series of RBD mutants in isogenic yeast-display assays, which recapitulated the deep mutational scanning measurements (Figures 4A-C), including confirmation that some mutations enhance expression (V367F and G502D) or ACE2 affinity (N501F, N501T, and Q498Y). We next analyzed binding of purified mammalian-expressed RBD protein to monomeric human ACE2, using RBDsfrom seven sarbecoviruses (SARS-CoV-2, SARS-CoV-1, WIV1, RaTG13, SHC014, ZXC21, and ZC45) immobilized on the surface of biosensors using biolayer interferometry. These 1:1 binding affinities on purified proteins parallel the measurements made using dimeric ACE2 with our yeast-displayed libraries, confirming the validity of the approach (Figures 4D, S5). Finally, we tested how some mutations affected spike-mediated entry of pseudotyped lentiviral particles into ACE2-expressing target cells (Figure 4E) (Crawford et al., 2020). The trends observed for entry by the spike-pseudotyped lentiviral particles generally confirmed the deep mutational scanning measurements: three of four mutations that we identified as detrimental for RBD expression or ACE2 binding greatly reduced pseudovirus entry, while a mutation that had little phenotypic effect in the deep mutational scanning did not affect viral entry. We also tested two ACE2 affinity-enhancing mutations and found that both increased pseudovirus entry. Note that this result with single-cycle pseudovirus does not necessarily imply that these mutations would increase growth of authentic SARS-CoV-2, since multi-cycle viral replication often involves tuning of receptor affinity to simultaneously optimize viral attachment and release (Callaway et al., 2018; Hensley et al., 2009; Lang et al., 2020). Taken together, these experiments help validate the accuracy and relevance of the deep mutational scanning measurements.

**Figure 4.**
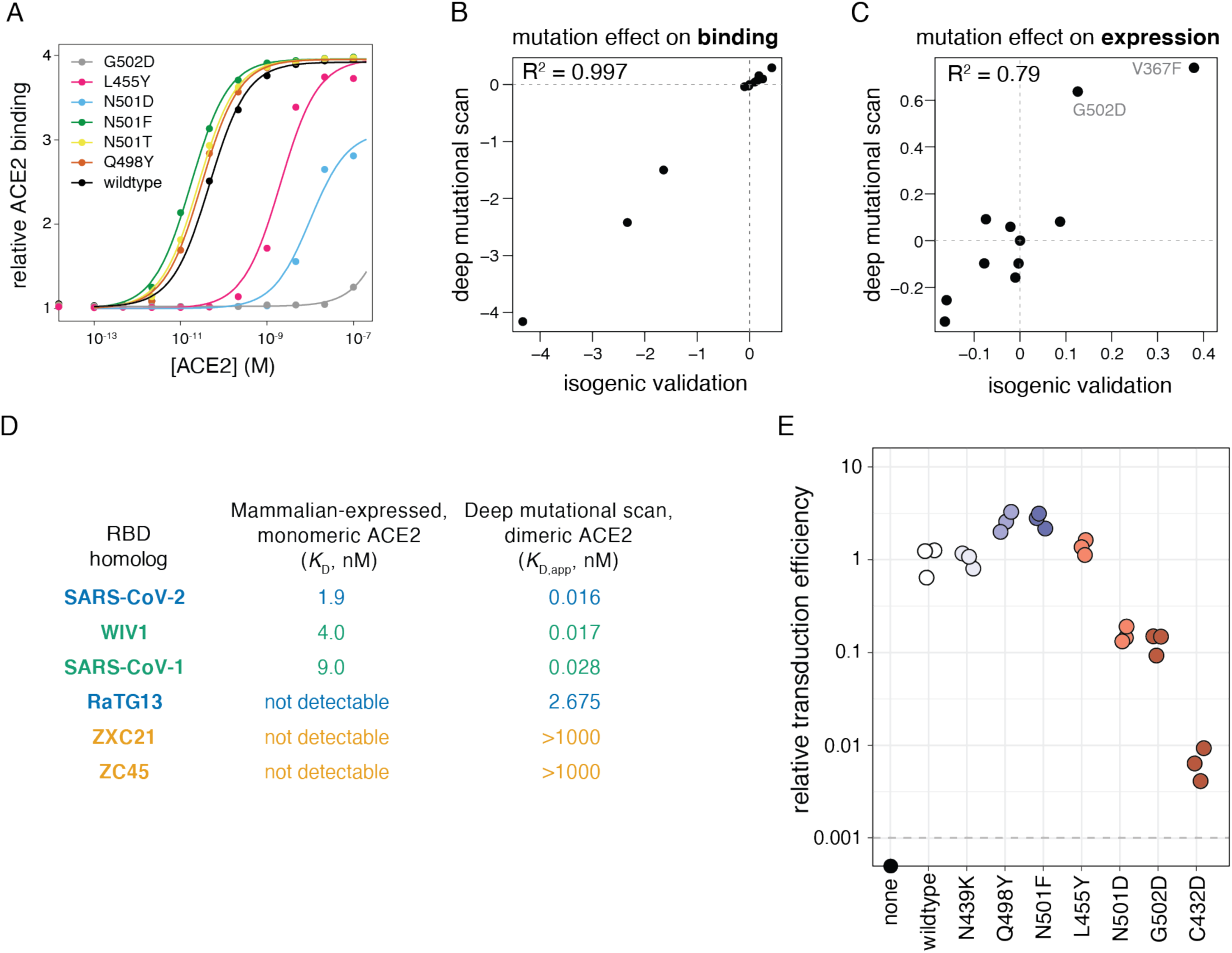
Validation of mutation effects measured in deep mutational scanning. (A) Titration curves for select mutations that were re-cloned and validated in isogenic cultures. (B, C) Correlation in binding (B) and expression (C) effects of mutations between deep mutational scanning and isogenic validation experiments, including mutants shown in (A) and Figure 8C. (D) Comparisons of dissociation constants measured for mammalian-expressed purified RBD binding to monomeric human ACE2 (see Figure S5) and yeast displayed RBD binding to natively dimeric ACE2 from our deep mutational scan. (E) Effects of mutations on transduction of ACE2-expressing cells by lentiviral particles pseudotyped with SARS-CoV-2 spike carrying the indicated mutation. Mutants are colored by their effects on ACE2 binding as measured in the deep mutational scanning using the same color scale as in Figure 3B (increased affinity in blue, reduced affinity in red). Titers that fell below the limit of detection (dashed horizontal line) are plotted on the x-axis.

### Interpreting mutation effects in the context of the RBD structure

To relate our sequence-phenotype maps to the RBD structure, we mapped the effects of mutations onto the ACE2-bound SARS-CoV-2 RBD crystal structure (Lan et al., 2020), coloring each residue’s C_α_ by the mean effect of amino-acid mutations at that site on expression (Figure 5A) or binding (Figure 5B). Interactive structure-based visualizations of specific residue sets discussed in the following sections can be found at the following link: https://jbloomlab.github.io/SARS-CoV-2-RBD_DMS/structures/.

**Figure 5.**
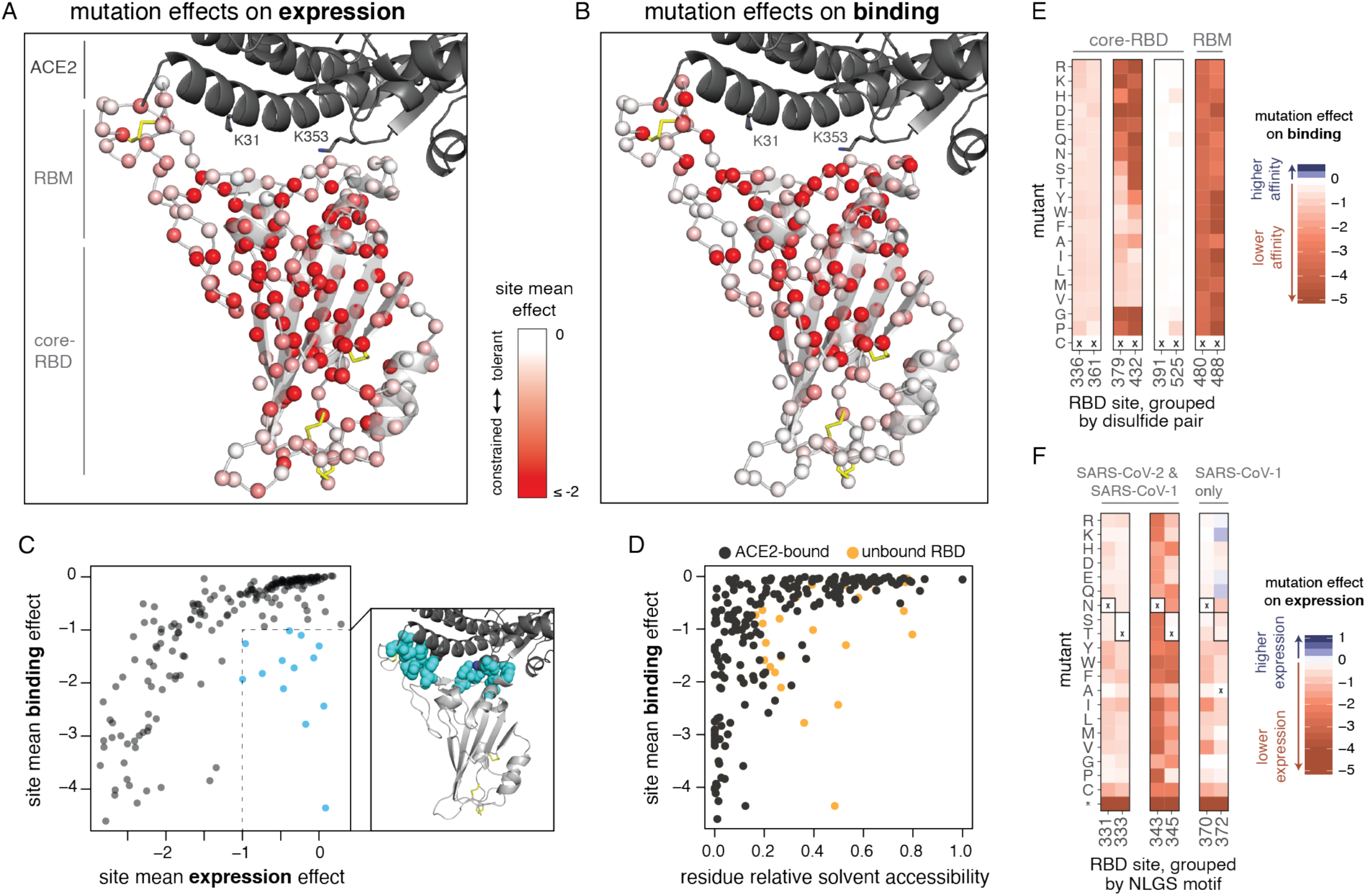
Mutation effects in the context of the RBD structure. (A, B) Mutational constraint mapped to the SARS-CoV-2 RBD structure. A sphere at each site’s C_α_ is colored according to the mean effect of amino-acid mutations at the site with respect to expression (A) or binding (B), with red indicating more constraint. RBD structural features and the ACE2 K31 and K353 interaction hotspot residues are labeled. Yellow sticks indicate disulfide bridges. Interactive structure-based visualizations of these data are at https://jbloomlab.github.io/SARS-CoV-2-RBD_DMS/structures/ (C) Relationship between mutational constraint on binding and expression. The structural view shows in cyan the sites that are under strong mutational constraint with respect to ACE2 binding but are tolerant of mutations with respect to expression. (D) Relationship between mutational constraint on binding and residue solvent accessibility (RSA). Black dots indicate RSA in the full ACE2-bound RBD structure, and when sites have large changes in RSA in the unbound structure, then their RSA in that structure is also shown in orange. (E) Mutation effects on binding at disulfide cysteine residues. Heatmaps as in Figure 3B. RBD sites are grouped by disulfide pair, and labeled according to location in the core-RBD or RBM sub-domains. (F) Mutation effects on expression at N-linked glycosylation sites (NLGS). RBD sites are grouped by NLGS motif (NxS/T, where x is any amino acid except proline). Boxed amino acids indicate those that encode a NLGS motif. NLGS motifs are labeled according to whether they are present in both the SARS-CoV-2 and SARS-CoV-1 RBD (N331 and N343 glycans), or in SARS-CoV-1 only (N370 glycan). See also Figure S6.

The two subdomains of the RBD differ in mutational constraint on expression and binding. The core-RBD subdomain consists of a central beta sheet flanked by alpha-helices, and presents a stably folded scaffold for the receptor binding motif (RBM, residues 437-508; (Li et al., 2005a)) which encodes ACE2-binding and receptor specificity (Letko et al., 2020). The RBM subdomain consists of a concave surface anchored by a β-hairpin and a disulfide bond stabilizing one of the lateral loops, which cradles the ACE2 α1 helix and a β-hairpin centered on K353_ACE2_. Consistent with the modularity of core-RBD-encoded stability and RBM-encoded binding, selection for expression primarily focuses on buried residues within the core-RBD (Figure 5A), while selection for binding focuses on the RBM-proximal core-RBD in addition to the RBM itself (Figure 5B), particularly on RBM residues that contact K31_ACE2_ and K353_ACE2_, which are “hotspots” of binding for SARS-CoV-1 and SARS-CoV-2 (Li, 2008; Shang et al., 2020; Wu et al., 2012).

Several ACE2-contact residues exhibit binding-stability tradeoffs, as has been seen in the active sites and binding interfaces of other proteins (Julian et al., 2017; Tokuriki et al., 2008; Wang et al., 2002). For example, several mutations to G502 enhance RBD expression (Figure 3A) but abolish binding (Figure 3B) due to steric clashes with ACE2 (Figure S6A). Similarly, mutations to polar amino acids enhance expression at interface residues Y449, L455, F486, Y505 (Figure 3A), consistent with the destabilizing effect of surface-exposed hydrophobic patches (Schwehm et al., 1998)—but these hydrophobic contacts form precise ACE2 packing contacts and are therefore required for binding (Figures 3B, S6B).

However, our data also indicate that global RBD stability contributes to ACE2-binding affinity. In general, mutation effects on RBD binding and expression are correlated (Figures 5C, S6C), with residues that deviate from this trend clustering at the ACE2 interface (Figure 5C, cyan points). This correlation between expression and binding is consistent with studies on antibodies, where mutations that improve stability and rigidity accompany increases in binding affinity (Davenport et al., 2016; Ovchinnikov et al., 2018; Schmidt et al., 2013). Because ACE2 binding is influenced by both global RBD stability and interface-specific constraints, a site’s tolerance to mutation is better explained by its extent of burial in the ACE2-bound RBD structure than its burial in the free RBD structure alone (Figure 5D).

Our data also reveal the importance of other specific sequence features. For example, the four disulfide bonds in the RBD have varying tolerance to mutation (Figures 5E, S6D), with the RBM C480:C488 disulfide completely constrained for ACE2 binding and C379:C432 being the most important of the core-RBD disulfides. The two RBD N-linked glycans contribute to RBD stability, as mutations that ablate the NxS/T glycosylation motif decrease RBD expression (Figure 5F). The SARS-CoV-1 RBD contains an additional RBD glycan, but its introduction at the homologous N370 in SARS-CoV-2 is mildly deleterious for expression (Figure 5F). However, there are other surface positions where introduction of NxS/T glycosylation motifs is tolerated or even beneficial (Figure S6E,F); adding glycans at some of these sites could be useful in resurfacing RBDs as antibody probes or epitope-focused immunogens (Duan et al., 2018; Eggink et al., 2014; Jardine et al., 2016; Kulp et al., 2017; Weidenbacher and Kim, 2019; Wu et al., 2010).

### Mutation effects at ACE2 contact sites and implications for sarbecovirus evolution

An initially surprising feature of SARS-CoV-2 was that its RBD tightly binds ACE2 despite differing in sequence from SARS-CoV-1 at many residues that had been defined as important for ACE2 binding by that virus (Andersen et al., 2020; Wan et al., 2020). Our map of mutational effects explains this observation by revealing remarkable degeneracy at ACE2 contact positions, with many mutations at interface sites being tolerated or even enhancing affinity (Figure 6A). Mutations that enhance affinity are particularly common at RBD sites Q493, Q498 and N501. Although these SARS-CoV-2 residues are involved in a dense network of polar contacts with ACE2 (Shang et al., 2020) (Figure 6B), our measurements show there is substantial plasticity in this network, as mutations that reduce the polar character of these residues often enhance affinity.

**Figure 6.**
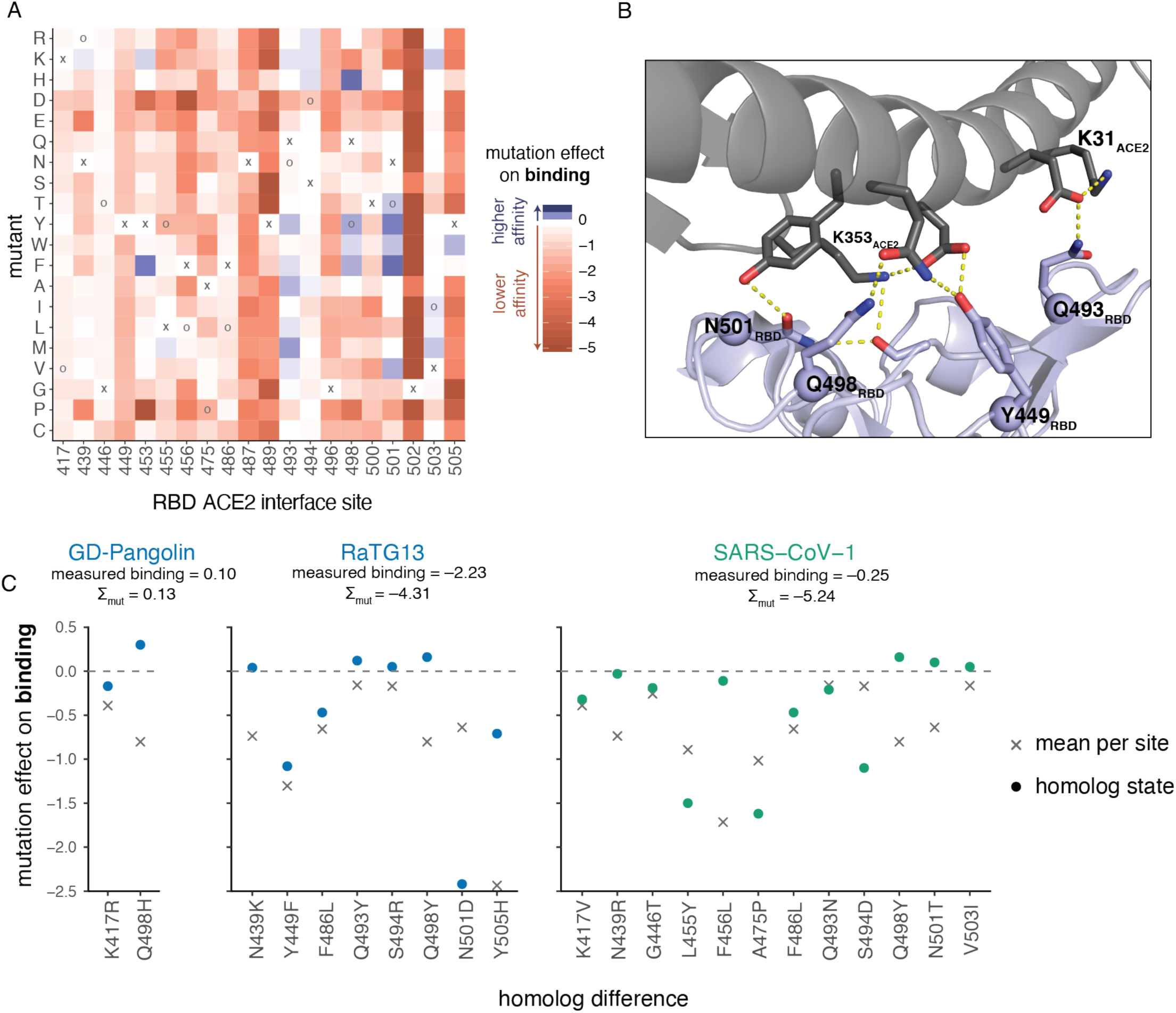
Mutation effects at ACE2 contact sites and implications for sarbecovirus evolution. (A) Heatmap as in Figure 3B, subsetted on sites that directly contact ACE2 in the SARS-CoV-2 or SARS-CoV-1 RBD structures, plus interface site 494 which is discussed as a key site of adaptation in the SARS-CoV-1 literature. (B) RBD sites Q493, Q498, and N501, which have many affinity-enhancing mutations, participate in polar contact networks involving the ACE2 interaction hotspot residues K31 and K353. (C) Variation at ACE2 contact sites in other sarbecovirus RBDs. The circles show the effects of the individual mutations that differentiate that virus’s ACE2 interface from SARS-CoV-2, while the x shows the mean effect of all mutations at that site. The sum of individual mutation effects at interface residues is shown, compared to the actual RBD binding relative to unmutated SARS-CoV-2.

Within the SARS-CoV-2 clade of sarbecoviruses (blue in Figure 1A), our maps of mutational effects on binding effectively explain variation in ACE2 affinity among different viruses. For example, GD-Pangolin has higher affinity for ACE2 than SARS-CoV-2 (Figures 1C, 2D), and this can be explained by the affinity-enhancing Q498H mutation present in this virus’s RBD sequence relative to SARS-CoV-2 (Figure 6C). In contrast, RaTG13 has substantially lower affinity for ACE2 than SARS-CoV-2 (Figures 1C, 2D), consistent with the presence of affinity-decreasing mutations including Y449F and N501D (Figure 6C). The fact that differences in binding affinity of GD-Pangolin and RaTG13 are well explained by summing the effects of individual mutations relative to SARS-CoV-2 suggests that our deep mutational scanning strategy is useful for sequence-based predictions of the ACE2-binding potential of future viruses isolated from the SARS-CoV-2 clade.

In contrast, the ACE2 binding interface of the more distantly related SARS-CoV-1 (Figure 1A) involves many more mutations relative to SARS-CoV-2, and this increased divergence appears to cause shifts in the actual effects of mutations on ACE2 binding. In particular, our deep mutational scanning shows that most of the SARS-CoV-1 amino-acid states are individually deleterious in SARS-CoV-2, despite being compatible with high-affinity binding in the SARS-CoV-1 background (Figure 6C). This shift in the effects of mutations between more distantly related RBDs is consistent with other studies of protein evolution demonstrating that epistastic entrenchment causes amino-acid preferences to change as homologous proteins become increasingly diverged (Hilton and Bloom, 2018; Lee et al., 2018; Pollock et al., 2012; Povolotskaya and Kondrashov, 2010; Shah et al., 2015; Starr and Thornton, 2016; Starr et al., 2018). Therefore, our current SARS-CoV-2 deep mutational scanning data are likely to be most useful for predicting the effects of mutations to RBDs closely related to that of SARS-CoV-2. However, the basic deep mutational scanning approach could easily be extended to also make measurements for other coronavirus clades.

### Mutational constraint of antibody epitopes

The RBD is the dominant target of neutralizing antibodies to SARS-CoV-2 (Brouwer et al., 2020; Cao et al., 2020; Ju et al., 2020; Premkumar et al., 2020; Rogers et al., 2020; Suthar et al., 2020; Yuan et al., 2020a; Zhang et al., 2020; Zost et al., 2020). It is unclear to what extent the RBD will evolve to escape such antibodies in a manner reminiscent of some other viruses (Smith et al., 2004; Trkola et al., 2005), although *in vitro* studies suggest that SARS-CoV-2 and SARS-CoV-1 RBDs are capable of fixing mutations that escape neutralizing antibodies (Baum et al., 2020; Rockx et al., 2010). To better define the RBD’s evolutionary capacity for antibody escape, we examined mutational constraint in the epitopes of antibodies with available structures that bind the SARS-CoV-1 or SARS-CoV-2 RBD (Figures 7A, S7A,B) (Hwang et al., 2006; Pak et al., 2009; Pinto et al., 2020; Prabakaran et al., 2006; Walls et al., 2019; Wrapp et al., 2020b; Wu et al., 2020; Yuan et al., 2020b).

**Figure 7.**
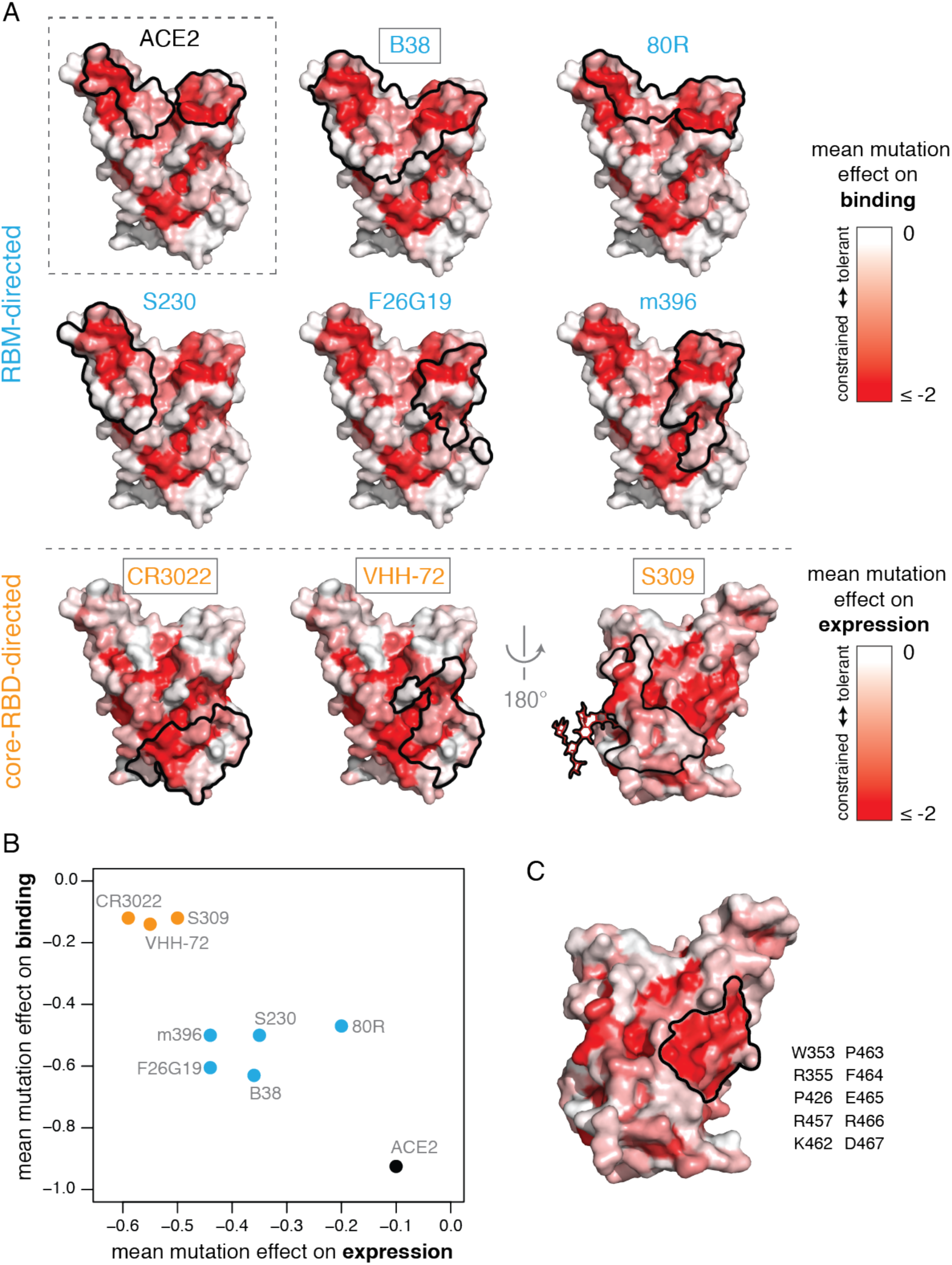
Mutational constraint of antibody epitopes. (A) Mutational constraint on surface residues comprising antibody epitopes. For each of 8 RBD-directed antibodies, black outlines indicate the epitope structural footprint, with surfaces colored by mutational constraint (red indicates more constrained). The direct ACE2 interface is shown in the upper-left, for reference. Names of antibodies capable of neutralizing SARS-CoV-2 are boxed. (Others neutralize SARS-CoV-1 but have not been demonstrated to neutralize SARS-CoV-2.) Constraint is illustrated as mutational effects on binding for RBM-directed antibodies (blue, top), and expression for core-RBD-directed antibodies (orange, bottom). The N343 glycan, which is present in the S309 epitope and is constrained with respect to expression, is shown only on this surface for clarity. (B) Average mutational constraint for binding and expression within each epitope. Points are colored according to the RBM versus core-RBD designation in (A). (C) Identification of a patch of mutational constraint surrounding RBD residue E465 which has not yet been targeted by any described antibodies. Surface is colored according to mutational effects on expression, as in (A, bottom). Residues in this constrained E465 patch are listed.

Many antibodies have epitopes that overlap the RBD ACE2 contact interface, and are therefore strongly constrained by mutation effects on binding (Figures 7A,B). For instance, antibodies B38 and 80R engage the two constrained patches that comprise the ACE2-binding interface, while S230, F26G19, and m396 engage either one of these ACE2-binding sub-regions. Of note, three additional antibodies have been reported that share a germline origin and structural epitope with B38, making the potential for viral escape from this “public” antibody of particular interest (Shi et al., 2020; Yuan et al., 2020a). However, none of the currently characterized antibodies have epitopes that are under as much functional constraint for binding as the ACE2-contact surface itself (Figure 7B), suggesting further epitope focusing could be achieved. The importance of such focusing is demonstrated by a recent study that identified RBD mutations enabling escape from RBM-directed neutralizing antibodies (Baum et al., 2020)—our data indicate that the escape occurs at sites that have high mutational tolerance (Figure S7C,D).

Epitopes of core-RBD-directed antibodies tend to be mutationally constrained with respect to expression rather than binding (Figures 7A,B). These core-RBD epitopes are highly conserved across the sarbecovirus alignment (Figure S7E), explaining the cross-reactivity of these antibodies between SARS-CoV-1 and SARS-CoV-2 (Huo et al., 2020; Pinto et al., 2020; Wrapp et al., 2020c). Although residues in these epitopes are constrained for stability even in our measurements on the isolated RBD, some of them likely exhibit additional constraint due to quaternary contacts made in the full spike trimer (Walls et al., 2020; Wrapp et al., 2020a; Yuan et al., 2020b). We identified an additional core-RBD patch centered on residue E465 that is also mutationally constrained (Figure 7C) and evolutionarily conserved (Figure S7E), but is not targeted by any currently known antibody and might represent a promising target for neutralization.

Taken together, our results identify multiple mutationally constrained patches on the RBD surface that can be targeted by antibodies. These findings provide a framework that could inform the formulation of antibody cocktails aiming to limit the emergence of viral escape mutants (Baum et al., 2020; Pinto et al., 2020; Wu et al., 2020; Zost et al., 2020), particularly if deep mutational scanning approaches like our own are extended to define RBD antibody epitopes in functional as well as structural terms (Dingens et al., 2019).

### Using sequence-phenotype maps to interpret genetic variation in SARS-CoV-2

An important question is whether any mutations that have appeared in circulating SARS-CoV-2 isolates have functional consequences. Despite intense interest in this question, experimental work to characterize the effects of SARS-CoV-2 mutations has lagged far behind their identification in viral sequences. Our comprehensive experimental maps of the phenotypic effects of mutations provide a direct way to interpret the impact of current and future genetic variation in the SARS-CoV-2 RBD.

To assess the phenotypic impacts of mutations that have appeared in the SARS-CoV-2 RBD to date, we downloaded all 31,570 spike sequences available from GISAID (Elbe and Buckland-Merrett, 2017) on May 27, 2020, and identified RBD amino-acid mutations present in high-quality clinical isolates. All observed RBD mutations are at low frequency, with 56 of the 98 observed mutations present only in a single GISAID sequence. The observed mutations are significantly less deleterious for both ACE2 binding and RBD expression than random single-nucleotide-accessible mutations (Figures 8A, S8A,B, P-value < 10^−6^ for binding and expression, permutation tests), consistent with the action of purifying selection. Purifying selection against deleterious mutations is especially apparent for mutations that are observed multiple times in circulating variants, with a substantial number of singletons being mildly or moderately deleterious whereas most mutations observed multiple times are largely neutral (Figures 8A, S8A,B). This general pattern of increased purifying selection on more common mutations is consistent with theoretical expectation and empirical patterns observed for other viruses (Pybus et al., 2007; Xue and Bloom, 2020).

**Figure 8.**
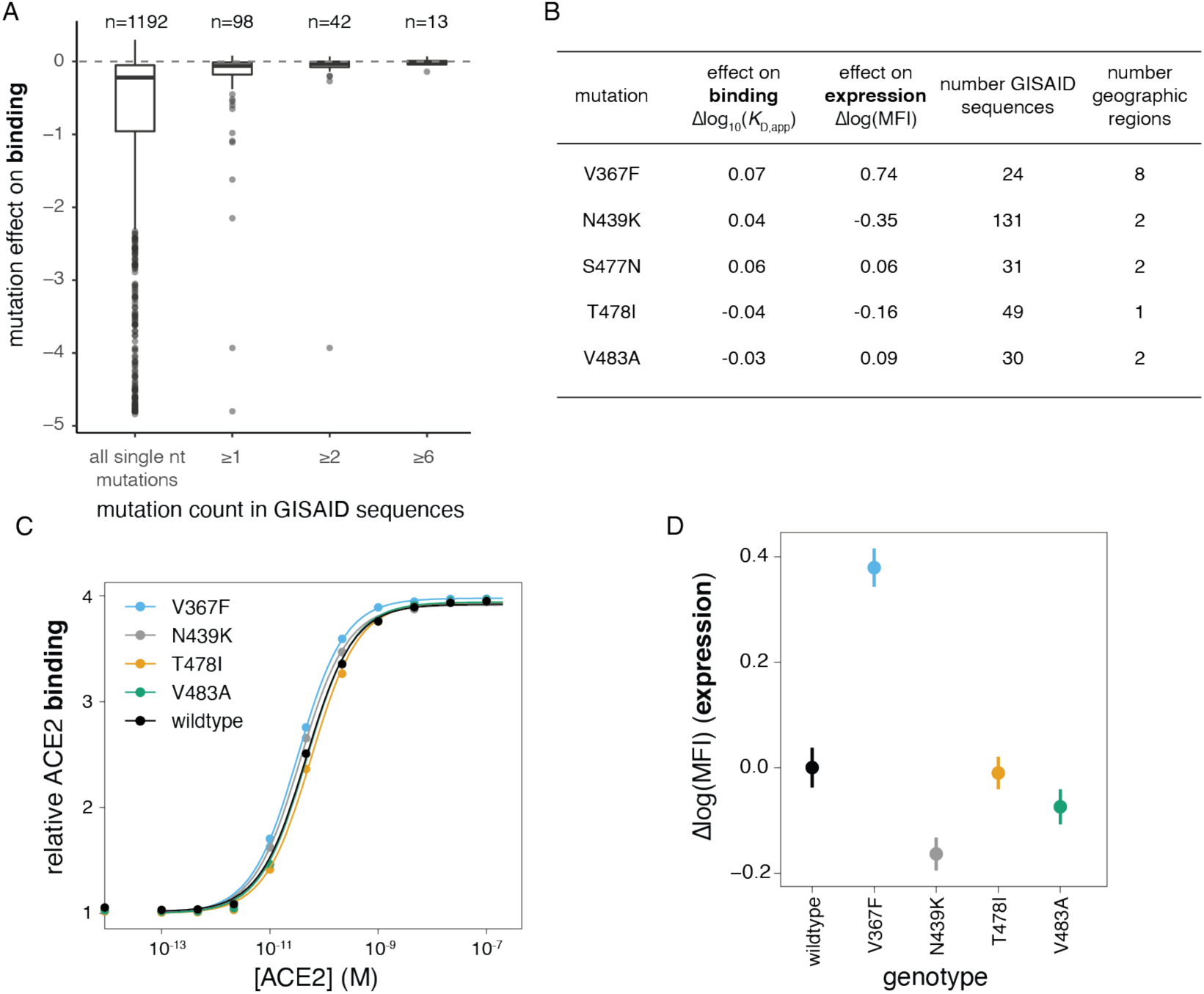
Phenotypic impacts of current genetic variation in the SARS-CoV-2 RBD. (A) Distribution of effects on ACE2 binding of mutations observed among circulating SARS-CoV-2 isolates. The distribution of mutation effects is shown for all amino-acid mutations accessible via single-nucleotide mutation from the SARS-CoV-2 Wuhan-Hu-1 gene sequence, compared to the distributions for subsets of mutations that are observed in sequenced SARS-CoV-2 isolates deposited in GISAID at increasing observation count thresholds. n, number of mutations in each subset. (B) Summary of most frequent mutations among GISAID sequences, reporting our deep mutational scanning measured effect on binding and expression, the number of GISAID sequences containing the mutation, and the number of geographic regions from which a mutation has been reported. (C, D) Validation of the mutational effects on binding (C) and expression (D) for 4 of the 5 most frequent circulating RBD variants. S477N rose to high frequency after we began our validation experiments, and so was not included. Error bars in (D) are standard error from 11 samples.

Our discovery of multiple strong affinity-enhancing mutations to the SARS-CoV-2 RBD raises the question of whether positive selection will favor such mutations, since the relationship between receptor affinity and fitness can be complex for viruses that are well-adapted to their hosts (Callaway et al., 2018; Hensley et al., 2009; Lang et al., 2020). Strong affinity-enhancing mutations are accessible via single-nucleotide mutation from SARS-CoV-2 (Figure S8C), but none are observed among circulating viral sequences in GISAID (Figure 8A), and there is no significant trend for actual observed mutations to enhance ACE2 affinity more than randomly drawn samples of all single nucleotide mutations (see permutation tests in Figure S8D). Taken together, we see no clear evidence of selection for stronger ACE2 binding, consistent with SARS-CoV-2 already possessing adequate ACE2 affinity at the beginning of the pandemic.

Last, we validated our deep mutational scanning measurements of the effects of mutations that are especially prevalent among naturally occurring sequences in GISAID. Our deep mutational scanning suggests small phenotypic effects for the most prevalent mutations, with the exception of V367F, which substantially enhances expression (Figure 8B). We re-cloned and tested most of these prevalent mutations for expression and ACE2 binding in isogenic yeast display assays. Consistent with the deep mutational scanning, the only large phenotypic effect was increased expression of V367F (Figure 8C,D). We also validated that N439K, the most prevalent RBD mutation which may have a very slight affinity-enhancing effect (Figures 8B,C), has no impact on entry of spike-pseudotyped lentiviral particles within the resolution of our assays (Figure 4E). The relevance of V367F’s stability-enhancing effect for viral fitness is unclear, though this mutation has independently arisen multiple times (van Dorp et al., 2020). Taken together, our results suggest that there is little phenotypic diversity in ACE2 binding among circulating variants at this early stage of the pandemic—although it will of course be interesting to use our maps to continually assess the phenotypic effects of future mutations as the virus evolves.

## Discussion

Vast numbers of viral genomes have been sequenced in almost real-time during the SARS-CoV-2 pandemic. These genomic sequences have been useful for understanding viral emergence and spread (Andersen et al., 2020; Bedford et al., 2020; Fauver et al., 2020), but the lack of corresponding high-throughput functional characterization means that speculation has greatly outpaced experimental data when it comes to understanding the phenotypic consequences of mutations. Here, we take a step toward providing phenotypic maps commensurate with the scale of genomic data by experimentally characterizing how all amino-acid mutations to the RBD affect the expression of folded protein and its affinity for ACE2, two key factors for viral fitness. These maps show that RBD mutations that have appeared in SARS-CoV-2 to date are slightly deleterious or nearly neutral with respect to these two biochemical phenotypes, with the exception of one mutation (V367F) that increases expression of folded protein. Notably, there has been no selection to date for any of the numerous evolutionarily accessible mutations that strongly enhance ACE2 binding affinity. The genetic diversity of SARS-CoV-2 is likely to increase as it continues to circulate in the human population, and so our phenotypic maps should become increasingly valuable for viral surveillance as mutations accumulate over time.

It is important to remember that our maps define biochemical phenotypes of the RBD, not how these phenotypes relate to viral fitness. There are many complexities in the relationship between biochemical phenotypes of yeast-displayed RBD and viral fitness. First, there are subtle differences in glycan structures between yeast versus human cells (Hamilton et al., 2003), though the overall role of glycans in RBD stability is preserved in yeast systems (Chen et al., 2014). Second, the RBD is just one domain of the viral spike, which engages in complex dynamic movements to mediate viral entry (Huo et al., 2020; Walls et al., 2019, 2020; Wrapp et al., 2020b). Finally, spike-mediated entry is just one component of fitness, which involves a myriad of incompletely understood factors that determine how well a virus spreads from one human to another (Kutter et al., 2018). To some degree, these caveats are universal of experimental studies, as even sophisticated animal models are imperfect proxies for true fitness (Louz et al., 2013)—but they are especially true for basic biochemical phenotypes like the ones we measure. However, on a hopeful note, our measurements correlate well with cellular entry by spike-pseudotyped viral particles expressing sarbecovirus RBD homologs (Figures 1D) and single mutants of the SARS-CoV-2 RBD (Figure 4E). Furthermore, fitness ultimately arises from the concerted action of biochemical phenotypes, which are in turn determined by genotype (Dean and Thornton, 2007; Harms and Thornton, 2013; Russell et al., 2014). By making the first link from mutations to biochemical phenotypes, we have taken a step towards enabling better interpretation of viral genetic variation.

One important area where our maps do have clear relevance is assessing the potential for SARS-CoV-2 to undergo antigenic drift by fixing mutations at sites targeted by antibodies, as occurs for some other viruses such as influenza (Smith et al., 2004). The RBD is the dominant target of neutralizing antibodies (Cao et al., 2020; Ju et al., 2020; Pinto et al., 2020; Rogers et al., 2020; Seydoux et al., 2020; Shi et al., 2020; Wu et al., 2020; Zost et al., 2020), and so any antigenic drift will be constrained by its mutational tolerance. Our results show that many mutations to the RBD are well-tolerated with respect to both protein folding and ACE2 binding. However, the ACE2 binding interface is more constrained than most of the RBD’s surface, which could limit viral escape from antibodies that target this interface (Rockx et al., 2010). In this respect, our maps enable several important observations. First, no characterized antibodies have epitopes that are as constrained as the actual RBD surface that contacts ACE2, suggesting that there is room for epitope focusing to minimize viral escape. Second, there are a number of RBD mutations that enhance ACE2 affinity, which implies ample evolutionary potential for compensation of deleterious mutations in the ACE2 interface in a manner reminiscent of multi-step escape pathways that have been described for other viruses (Bloom et al., 2010; Friedrich et al., 2004; Gong et al., 2013; Lynch et al., 2015; Wu et al., 2017). It should be possible to shed further experimental light on the potential for antigenic drift by extending our deep mutational scanning methodology to directly map immune-escape mutations as has been done for other viruses (Dingens et al., 2019; Lee et al., 2019; Wu et al., 2019).

RBD-based antigens represent a promising vaccine approach (Chen et al., 2020a, 2020b; Quinlan et al., 2020; Ravichandran et al., 2020; Zang et al., 2020). Our sequence-phenotype maps can directly inform efforts to engineer such vaccines in several ways. First, we identify many mutations that enhance RBD expression, a correlate of thermodynamic stability (Kowalski et al., 1998a, 1998b; Shusta et al., 1999) and a desirable property in vaccine immunogens. Second, our maps show which mutations can be introduced into the RBD without disrupting key biochemical phenotypes, thereby opening the door to resurfacing immunogens to focus antibodies on specific epitopes (Duan et al., 2018; Eggink et al., 2014; Jardine et al., 2016; Kulp et al., 2017; Weidenbacher and Kim, 2019; Wu et al., 2010). Finally, our maps show which surfaces of the RBD are under especially strong constraint and might thereby be targeted by structure-guided vaccines to stimulate neutralizing immunity with breadth across the sarbecovirus clade: in addition to the ACE2 interface itself, these surfaces include several core-RBD surface patches targeted by currently described antibodies and a previously undescribed core-RBD surface patch surrounding residue E465.

Finally, our work should be useful for understanding the evolution of sarbecoviruses more broadly, including the potential for more spillovers into the human population. There is a dizzying diversity of RBD genotypes and phenotypes among sarbecoviruses within bat reservoirs (Boni et al.; Demogines et al., 2012; Frank et al., 2020; Hu et al., 2017; Latinne et al., 2020; Letko et al., 2020; MacLean et al., 2020). A prerequisite for these viruses to jump to humans is the ability to efficiently bind human receptors (Becker et al., 2008; Letko et al., 2020; Menachery et al., 2015, 2016). Our maps are immediately useful in assessing the effects on ACE2-binding of mutations to viruses within the SARS-CoV-2 clade, and extensions to account for epistasis and genetic background could further inform understanding of the evolutionary trajectories that enable sarbecoviruses to efficiently infect human cells.

## Acknowledgements

We thank Keara Malone for experimental assistance, Adam Dingens, Katherine Xue, Dan Ellis, and Neil King for helpful suggestions, and Frederick Matsen for intellectual support and hospitality while these experiments were being carried out. We thank the Flow Cytometry and Genomics core facilities at the Fred Hutchinson Cancer Research Center for experimental support. This work was supported by the NIAID / NIH (R01AI141707 and R01AI12893 to J.D.B., HHSN272201700059C to D.V., F30AI149928 to K.H.D.C., and T32AI083203 to A.J.G.), NIGMS / NIH (R01GM120553 to D.V.), a Pew Biomedical

Scholars Award to D.V., Burroughs Wellcome Investigators in the Pathogenesis of Infectious Diseases awards to D.V. and J.D.B., the Bill & Melinda Gates Foundation (OPP1156262 to D.V.), and Fast Grants (to D.V.). T.N.S. is a Washington Research Foundation Innovation Fellow at the University of Washington Institute for Protein Design and a Howard Hughes Medical Institute Fellow of the Damon Runyon Cancer Research Foundation. J.D.B. is an Investigator of the Howard Hughes Medical Institute.

## Author contributions

Conceptualization, T.N.S., D.V., and J.D.B.; Methodology, T.N.S. and J.D.B.; Investigation, T.N.S. and A.J.G.; Code, T.N.S., S.K.H., K.H.D.C., and J.D.B.; Formal Analysis, T.N.S. and J.D.B.; Validation, A.J.G., K.H.D.C., M.J.N., J.E.B., M.A.T., and A.C.W.; Visualization, T.N.S., S.K.H., and J.D.B.; Writing – original draft, T.N.S. and J.D.B.; Writing – review and editing, T.N.S., A.J.G., S.K.H., K.H.D.C., D.V., J.D.B.; Supervision, D.V. and J.D.B.

## Methods

### Data and Code Availability

We provide all data and code in the following ways:

- Raw data tables of our replicate functional scores at the level of single mutations (Supplemental File 3, and GitHub:https://github.com/jbloomlab/SARS-CoV-2-RBD_DMS/blob/master/results/single_mut_effects/single_mut_effects.csv)
- Raw data tables of our replicate functional scores among sarbecovirus homologs (Supplemental File 1 and GitHub:https://github.com/jbloomlab/SARS-CoV-2-RBD_DMS/blob/master/results/single_mut_effects/homolog_effects.csv)
- Illumina sequencing counts for each barcode among FACS bins (https://github.com/jbloomlab/SARS-CoV-2-RBD_DMS/blob/master/results/counts/variant_counts.csv)
- The complete variant:barcode lookup table (https://github.com/jbloomlab/SARS-CoV-2-RBD_DMS/blob/master/results/variants/codon_variant_table.csv)
- The complete computational workflow to generate and analyze these data, including reproducible code within a programmatically constructed computational environment (https://github.com/jbloomlab/SARS-CoV-2-RBD_DMS)
- A Markdown summary of the organization of analysis steps, with links to key data files and Markdown summaries of each step in the analysis pipeline (https://github.com/jbloomlab/SARS-CoV-2-RBD_DMS/blob/master/results/summary/summary.md), with specific Markdown summaries linked in the relevant Methods sections below
- All raw sequencing data are uploaded to the NCBI Short Read Archive (BioProject PRJNA639956).

### RBD cloning

The Spike receptor binding domain (RBD) from SARS-CoV-2 (isolate Wuhan-Hu-1, Genbank accession number MN908947, residues N331-T531) and additional sarbecovirus homologs (RaTG13, Genbank MN996532; GD-Pangolin consensus from (Lam et al., 2020); SARS-CoV-1 Urbani, Genbank AY278741; WIV1, Genbank KF367457 (identical RBD sequence to WIV16); LYRa11, Genbank KF569996; Rp3, Genbank DQ071615; HKU3-1, Genbank DQ022305; Rf1, Genbank DQ412042; ZXC21, Genbank MG772934; ZC45, Genbank MG772933; and BM48-31, Genbank NC014470) were ordered as yeast codon-optimized gBlocks (IDT) and cloned into the pETcon yeast surface display expression vector. The destination vector was modified downstream from the yeast surface display fusion construct to include a barcode landing pad for subsequent library generation, along with Illumina sequencing priming handles for downstream barcode sequencing and NotI digestion sites for downstream PacBio sequencing preparation. This plasmid sequence is provided on GitHub at https://github.com/jbloomlab/SARS-CoV-2-RBD_DMS/tree/master/data/plasmid_maps/2649_pETcon-SARS-CoV-2-RBD-201aa.gb.

### Isogenic yeast display induction and titration

RBD variant plasmids were transformed into the AWY101 *Saccharomyces cerevisiae* strain (Wentz and Shusta, 2007), selecting for the plasmid Trp auxotrophic marker on SD-CAA selective plates (6.7g/L Yeast Nitrogen Base, 5.0g/L Casamino acids, 1.065g/L MES acid, and 2% w/v dextrose). Single colonies were inoculated into 1.5mL liquid SD-CAA media, and grown overnight at 30°C. Then 1 OD unit of yeast were back-diluted into 1.5mL SG-CAA+0.1%D induction media (2% w/v galactose supplemented with 0.1% dextrose), and incubated for 16-18 hours at room temperature.

Induced cells were spun down at 250,000 cells per sample and washed in PBS-BSA (0.2mg/mL). Samples were resuspended in primary labeling solutions across a range of concentrations of biotinylated human ACE2 ectodomain (Figure 1C), which contains its natural dimerization domain (ACROBiosystems AC2-H82E6). Primary labeling reactions were conducted in sufficient reaction volumes for each concentration to avoid ligand depletion effects of greater than 10%. For instance, the lowest sample concentration of 10^−13^ M was scaled to 25mL, at which volume 2.9% of total ligand molecules are estimated to be titrated in RBD:ACE2 complexes given the wildtype *K*_D,app_ and an estimated 50,000 surface RBDs per cell (Boder and Wittrup, 1997). Following overnight equilibration of ACE2 binding at room temperature, cells were washed in ice-cold PBS-BSA, and resuspended in PBS-BSA containing 1:200 diluted FITC-conjugated anti c-Myc antibody (Immunology Consultants Lab, CMYC-45F) to label for RBD surface expression via a C-terminal c-Myc epitope tag, and 1:200 diluted PE-conjugated streptavidin (Thermo Fisher S866) to detect bound biotinylated ACE2 ligand. Following 1 hour of secondary labeling at 4°C, cells were washed twice in ice-cold PBS-BSA, and resuspended in PBS.

RBD surface expression and ACE2-binding levels were determined via flow cytometry using a BD LSRFortessa X-50. For the flow cytometry, 10,000 cells were analyzed at each ACE2 concentration across a titration series. Cells were gated to select for singleton events, FITC labeling was used to subset RBD+ cells, and PE labeling was measured within this FITC+ population. To mimic the subsequent library sorting experiments in which we are blinded to exact PE fluorescence within a given PE fluorescence bin (since we only sequence barcodes within a bin), we analyzed isogenic titration data by drawing equivalent bins of PE fluorescence that capture 95% of unbound unmutated SARS-CoV-2 cells (bin1), 95% of saturated SARS-CoV-2 cells (bin4), and a bin2/bin3 boundary evenly spaced on the log-scale between the boundaries of the bin1 and bin4 partitions (see Figure 2B). For each ACE2 concentration, we determine the mean bin of PE fluorescence as a simple weighted mean value across integer-weighted bins:

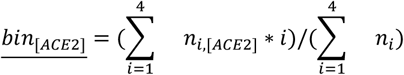

where *n*_*i,[ACE2]*_ is the number of cells that fall into bin *i* at a given ACE2 concentration, and *i* is the simple integer value of a bin from 1 to 4.

We determined the binding constant *K*_D,app_ describing the affinity of each RBD variant for human ACE2 ligand along with free parameters *a* (titration response range) and *b* (titration curve baseline) via nonlinear least squares regression using a standard non-cooperative Hill equation relating the mean bin response variable to the ACE2 labeling concentration:

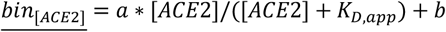

We report apparent *K*_D_ values (*K*_D,app_) that do not take into account the stoichiometry of the multivalent yeast-displayed RBD interaction with dimeric ACE2. Following this “apparent” nomenclature, we report ACE2 concentrations as molarity of the monomeric subunit. A computational notebook detailing the fits of all isogenic RBD titrations is provided on GitHub (https://github.com/jbloomlab/SARS-CoV-2-RBD_DMS/blob/master/data/isogenic_titrations/homolog_validations.md and https://github.com/jbloomlab/SARS-CoV-2-RBD_DMS/blob/master/data/isogenic_titrations/point-mut-validations.md).

### Library mutagenesis

Mutagenesis of the SARS-CoV-2 RBD was performed in two independent replicates via the method described in (Bloom, 2014) with the modification that primers lengths were adjusted to ensure equal melting temperatures as described in (Dingens et al., 2017) and we used NNS rather than NNN primers. Our general library generation and sequencing workflow is outlined in Figure S1A. Briefly, we designed nested mutagenic primers containing degenerate NNS codons that tile across the SARS-CoV-2 RBD, which were ordered as oPools from Integrated DNA Technologies. The script used to design the mutagenic primers and the resulting primers are available at https://github.com/jbloomlab/SARS-CoV-2-RBD_DMS/tree/master/data/primers/mutational_lib. We conducted three rounds of mutagenesis, each consisting of 7 mutagenic PCR cycles and 20 joining PCR cycles. The final joined products were amplified for 10 cycles with primers that append a unique identifier N16 barcode sequence to the 3’ end of each mutagenized insert, downstream from the RBD stop codon and mRNA 3’ UTR. Barcodes were also PCR appended to the un-mutagenized RBD homologs via the same primer addition PCR. Primers used in library assembly are provided on GitHub (https://github.com/jbloomlab/SARS-CoV-2-RBD_DMS/tree/master/data/primers).

Mutagenized SARS-CoV-2 libraries and pooled wildtype homolog RBDs were cloned into EcoRI-HF/SacI-HF digested pETcon vector (sequence linked above) using NEBuilder HiFi DNA Assembly (NEB E2621). Assembled products were Ampure purified and electroporated into electrocompetent NEB10-beta cells. Electroporated cells were plated on 15cm LB+ampicillin plates at an estimated bottleneck of 100,000 (SARS-CoV-2 mutant libraries) or 1,000 (pooled RBD homologs) colony forming units to limit library size. After approximately 18 hours of outgrowth, colonies were scraped into liquid LB+ampicillin, and grown for 2.5 hours in liquid culture prior to plasmid purification.

Plasmid pools were transformed into the AWY101 strain of *Saccharomyces cerevisiae* via the protocol of Gietz and Schiestl (Gietz and Schiestl, 2007). SARS-CoV-2 mutant libraries were transformed at 50ug scale and the pooled RBD homolog controls were transformed at 10ug scale. Colony forming unit counts from plated serial dilutions indicate transformation yield of >1 million cfus. Transformed yeast grew for 14 hours post-transformation in 100mL selective SD-CAA media, and were subsequently back-diluted into 100mL fresh SD-CAA at 1 OD600 for an additional 9 hours passage, to enable further resolution of multiple vector transformants (Scanlon et al., 2009). Transformed yeast libraries were flash frozen in 1e8 cfu aliquots and stored -80°C.

### PacBio library sequencing and analysis

PacBio sequencing was used to acquire long sequence reads spanning the N16 barcode and the RBD gene sequence. PacBio sequencing inserts were prepared from bacterially-purified plasmid pools via NotI-HF restriction digest followed by gel purification and SMRTbell ligation. The use of restriction digest rather than PCR eliminates the possibility of PCR strand exchange scrambling barcodes. Each SARS-CoV-2 RBD mutant library was spiked to 1% frequency with the internal standard pool of RBD homologs. Each replicate library was sequenced in two SMRT Cells on a PacBio Sequel using 20-hour movie collection times. PacBio circular consensus sequences (CCSs) were generated from the raw subreads using the `ccs` program (https://github.com/PacificBiosciences/ccs, version 4.2.0), setting the parameters to require 99.9% accuracy and a minimum of 3 passes. The resulting CCSs are available on the NCBI Sequence Read Archive at https://www.ncbi.nlm.nih.gov/bioproject/PRJNA639956.

We then processed the CCSs to identify the RBD sequence (SARS-CoV-2 or one of the 11 homologs), call any mutations in the RBD sequence, and determine the associated 16-nucleotide barcode. To do this, we used `alignparse` (Crawford and Bloom, 2019), which in turn makes use of `minimap2` (Li, 2018), version 2.17. We only retained CCSs that matched the parental RBD sequence with no more than 45 nucleotide mutations (corresponding to up to 15 codon mutations), had a barcode of the expected 16 nucleotide length, and had no more than one mismatch in the flanking regions expected in the sequenced amplicon. A computational notebook providing full details is available on GitHub at https://github.com/jbloomlab/SARS-CoV-2-RBD_DMS/blob/master/results/summary/process_ccs.md.

We next used these processed CCSs to generate a codon-variant lookup table that links each barcode to its associated codon mutations in the RBD sequence. To do this, we first filtered only for CCSs where the PacBio `ccs`-reported accuracy was at least 99.99% in both the RBD gene sequence and the barcode (the vast majority of CCSs passed this filter). We then determined the empirical accuracy of the CCSs by determining the concordance between the RBD gene sequence called by CCSs with the same barcode using the method implemented at https://jbloomlab.github.io/alignparse/alignparse.consensus.html#alignparse.consensus.empirical_accuracy. For both libraries, the empirical accuracy of the entire region of the CCS covering the RBD sequence was 99.8% if we ignored those with indels (Figure S1E). Most barcodes were covered by multiple CCSs (Figure S1E), and in that case we built a consensus of these CCSs after discarding any barcodes for which the CCSs differed often or at many sites using the method implemented at https://jbloomlab.github.io/alignparse/alignparse.consensus.html#alignparse.consensus.simple_mutconsensus. Finally, we discarded any variants with indels in the RBD. Therefore, more than 99.8% of the final barcode-linked variants should have the correctly determined RBD sequence, since 99.8% is the accuracy for those covered by just one CCS and most variants were called by the consensus of multiple CCSs. For further analysis of the barcoded variants, we then created a codon variant table using `dms_variants` (https://jbloomlab.github.io/dms_variants/, version 0.6.0). The final barcode-variant lookup table (which associates each barcode with its RBD sequence) is at https://github.com/jbloomlab/SARS-CoV-2-RBD_DMS/raw/master/results/variants/codon_variant_table.csv. Some summary statistics about the final composition of the libraries are in Figure S1, and the complete code used to generate the barcode-variant lookup table and many additional plots characterizing the composition of the libraries are on GitHub at https://github.com/jbloomlab/SARS-CoV-2-RBD_DMS/blob/master/results/summary/build_variants.md.

### Deep mutational scanning library yeast surface-display induction and labeling

Yeast libraries were thawed and grown overnight at 30°C in 180mL SD-CAA media at an initial OD600 of 0.1. We spiked our SARS-CoV-2 mutant libraries with the barcoded RBD homolog pool at a total fraction of 0.6% yeast density, such that each RBD homolog barcode should be present at a frequency on the same order of magnitude as the typical SARS-CoV-2 variant barcode. To induce RBD surface expression, yeast were back-diluted to 50mL (expression experiments) or 200mL (binding experiments) SG-CAA+0.1%D induction media at 0.67 OD600 and incubated at room temperature for 16-18 hours with mild agitation. Induced cells were spun and washed twice with PBS-BSA, before proceeding with ligand labeling.

For library expression experiments, 45 OD units yeast were washed twice with PBS-BSA and labeled in 3mL 1:100 diluted anti-Myc-FITC antibody for 1hr at 4°C with gentle mixing. Labeled cells were washed twice in PBS-BSA and resuspended in 5mL PBS for FACS. For library binding experiments, 8 OD units yeast per titration concentration (10^−13^ M to 10^−6^ M ACE2 at half-log intervals, plus a 0M ACE2 sample) were washed with PBS-BSA, and incubated with ACE2 ligand overnight at room temperature with gentle agitation. Labeling volumes were scaled at low ACE2 concentration to limit ligand depletion effects, as with isogenic titrations described above. Following equilibration of ACE2 labeling, cells were kept chilled while washing once with PBS-BSA, labeling for one hour in 1mL PBS-BSA with 1:100 diluted Myc-FITC and 1:200 Streptavidin-PE, washed two more times with PBS-BSA, and resuspended to 1mL in PBS.

### Fluorescence activated cell sorting (FACS) of yeast libraries

Yeast libraries were sorted into bins of FITC or PE fluorescence using a BD FACS Aria II. Cells were sorted into 5mL FACS tubes containing 1mL of 2xYPAD supplemented with 1% BSA. Tubes were pre-wet with collection media prior to sample collection, to reduce sticking and improve post-sort yield.

For expression sorts, cells were gated for singleton events (Figure S2A), followed by partitioning into four bins of FITC fluorescence (Figures 2A): bin 1 captures 99% of unstained cells, and bins 2-4 split the remaining library fraction into tertiles. We sorted >50 million cells from each library into these bins. From these same inductions, we also sorted 15 million RBD+ cells from each library (P4 population, Figure S2A), to enrich RBD-expressing cells within our libraries for our titration sorting experiments.

For ACE2-binding titrations, we gated cells for singleton events and RBD+ expression (Figure S2B). For each ACE2 concentration sample, we sorted cells into four bins of PE fluorescence as described above: bin1 captures 95% of unmutated SARS-CoV-2 cells incubated with 0M ACE2, bin4 captures 95% of unmutated cells at saturating ACE2 ligand, and the bin2/bin3 boundary evenly splits the log-MFI scale between the bin1 and bin4 boundaries (Figure 2B). We sorted each ACE2 concentration sample into these four bins for approximately 15 minutes, capturing 5-6 million cells per ACE2 concentration.

Following each sort, cells from each collection tube were spun for 5 min at 3,000 g in a tabletop centrifuge, yielding a visible pellet for any sample with at least ∼500,000 collected cells. Collection supernatant was removed, and cells were resuspended in SD-CAA media supplemented with 1:100 penicillin-streptomycin. Cells were resuspended to an estimated 2e6 cells/mL in 15mL culture tubes or baffled flasks for expresion post-sort samples, 5e5 cells/mL in baffled flasks for RBD+ sort samples, and 1mL (<1e6 cells) or 1.5mL (>1e6 cells) in 96-deep-well plates for titration samples. For expression FACS experiments, total cell recovery from all samples was measured via serial dilution and plating on YPD and SD-CAA plates for each sample, which showed average cellular recovery of 85% (range 79-94%), with 62% (range 52-77%) of cells retaining plasmid, with exception of the FITC-negative bin 1 populations, which showed 20% plasmid retention. These per-sample cell recovery counts were used to calibrate downstream sequencing numbers for the actual number of cells that grew out from each sort bin. For titration sorts, we did not titer all 64 post-sort samples, but instead spot checked 6 samples to ensure normal levels of cell recovery, which showed an average 66% cell recovery and 46% plasmid retention. As we did not titer all samples, we use the FACS log cell count as the estimate of number of cells collected in each bin, which makes the assumption that there are no systematic differences in post-sort cell yield across bins, which is more appropriate for these titration sorts where the ACE2 binding gates are nested within an overall RBD+ selection gate that selects for even plasmid retention (Figure S2B).

Post-sort samples were grown overnight in liquid media at 30°C. Plasmids were purified from post-sort yeast samples of <4e7 cfu using Zymo Yeast Miniprep kits (single column or 96-well plate formats) according to kit instructions, but with the addition of >2 hours Zymolyase treatment and a -80°C freeze/thaw cycle prior to cell lysis.

### Illumina Sequencing

Post-sort plasmid samples were PCR amplified from 10uL plasmid template input using primers flanking the N16 barcode that append remaining Illumina sequencing handles that are not already plasmid encoded, and unique NextFlex sample indices (https://github.com/jbloomlab/SARS-CoV-2-RBD_DMS/tree/master/data/primers). PCRs were conducted with KOD polymerase for 20 cycles, except for titration sort samples of less than 10,000 cells, where 28 cycles were necessary to obtain sufficient PCR product due to low sample input:

1. 95°C, 2min
2. 95°C, 20s
3. 58°C, 10s
4. 70°C, 10s
5. Return to 2, 19x (27x for low-input samples)

PCR products were Ampure purified, quantified via PicoGreen, and pooled to mirror desired sample frequencies given cell counts in each FACS sample. Pooled samples were gel purified, Ampure purified, and submitted for 2 lanes of 50bp single end Illumina HiSeq sequencing per library.

Demultiplexed reads were aligned to library barcodes determined from PacBio sequencing, yielding a count of the number of times each library barcode was sequenced within each FACS partition. Read counts for each FACS sample were downweighted by the ratio of total reads from a bin compared to the number of cells that were actually sorted into that bin. For one bin in which the number of HiSeq reads was less than the number of cells sorted into a bin, we re-amplified PCR product from a newly purified plasmid aliquot, and obtained reads via a single lane of MiSeq 50bp single end sequencing. Computational notebooks providing additional details on our Illumina sequencing processing and statistics are provided on GitHub (https://github.com/jbloomlab/SARS-CoV-2-RBD_DMS/blob/master/results/summary/count_variants.md and https://github.com/jbloomlab/SARS-CoV-2-RBD_DMS/blob/master/results/summary/analyze_counts.md).

### Calculating variant phenotypes for expression

For each library variant, we estimated mean expression based on its distribution of cell counts across FITC sort bins and the known censored fluorescence boundaries of each sort bin using a maximum likelihood approach (Peterman and Levine, 2016), enacted in the fitdistrplus R package (Delignette-Muller and Dutang, 2015), assuming the uncensored log-transformed fluorescence values for a genotype follow a normal distribution. Expression measurements were retained for barcodes for which at least 20 cells were sampled across the four sort bins, resulting in measured expression phenotypes for 92.9 and 90.5% of variants in libraries 1 and 2, respectively.

Expression measurements were represented as the difference in log-mean fluorescence intensity (MFI) relative to wildtype (ΔlogMFI = logMFI_variant_ - logMFI_wildtype_), such that a positive value indicates higher RBD expression. A very small fraction of wildtype and synonymous barcodes were ascribed non-fluorescing phenotypes, likely reflecting expression-abolishing mutations that occurred outside of the PacBio sequencing window. These variants were selected out prior to titration measurements by our RBD+ pre-sort, but remain in the expression measurements. To avoid artificially depressing the wildtype SARS-CoV-2 expression measurement and therefore miscalibrating this Δlog(MFI) scale, potentially annotating slightly deleterious mutational effects as beneficial, we computed the mean wildtype expression excluding these outliers (logMFI < 10.2 or 10.1 in lib1 and lib2, respectively). We note that we are unable to do the same for any library mutants for which we observe non-fluorescence, because we are unable to *a priori* determine whether a lack of expression is due to the library mutation versus external, unobserved factors. This uncertainty makes our calling of expression-enhancing mutations conservative, as mutational effects, if biased by these outliers, will tend to be pulled slightly down in their measurement. The global epistasis approach we explain below can mitigate the influence of these outlier observations on our final estimates of mutational effects. A computational notebook presenting our calculation of expression phenotypes and results is included on GitHub (https://github.com/jbloomlab/SARS-CoV-2-RBD_DMS/blob/master/results/summary/compute_expression_meanF.md).

### Calculating variant phenotypes for ACE2-binding affinity

For each library barcode at each ACE2 sample concentration, we determined its simple mean bin of ACE2-binding via the equation used above in isogenic titrations. We fit titration curves as above to determine barcode-specific *K*_D,app_ from the series of FACS-seq derived mean bin measurements across ACE2 concentration (Figure S2C). Because a barcode’s mean bin might be measured with varying certainty across different bins, we used weighted least squares nonlinear regression, weighing each mean bin estimate by an empirical variability estimate based on the per-sample cell count, derived from estimates of variability in repeated wildtype/synonymous barcode measurements grouped by sampling depth. To avoid fits of errant titration curves, we constrained the baseline parameter *b* to be fit between 1 and 1.5, and the response parameter *a* to be fit between 1.5 and 3. Through initial curve fit constraints and subsequent QC filtering, our fit *K*_D,app_ binding constants were constrained to be within the concentration range of our titration (10^−13^ – 10^−6^ M), and therefore many barcodes are censored at the upper limit with true *K*_D,app_ ≥ 10^−6^ M. We filtered out titration curves fit for variants with an average cell count <5 across sample concentrations, or with cell count <2 in 7 or more of the 16 samples. Finally, we filtered out the 5% of curves with the highest normalized mean square residual, where residuals are normalized from 0 to 1 by the fit response parameter *a*, such that titration curves that plateau at lower levels of saturated binding don’t have systematically smaller mean square residuals. This process yielded *K*_D,app_ estimates for 75.2 and 75.4% of variants in libraries 1 and 2, respectively. Binding measurements were represented as the difference in log_10_(*K*_D,app_) relative to wildtype (Δlog_10_(*K*_D,app_) = log_10_(*K*_D,app_)_wildtype_ – log_10_(*K*_D,app_)_variant_), polarized such that a positive value indicates higher variant ACE2 affinity. A computational notebook presenting our calculation of expression phenotypes and results is included on GitHub (https://github.com/jbloomlab/SARS-CoV-2-RBD_DMS/blob/master/results/summary/compute_binding_Kd.md).

### Decomposing single-mutant effects from multiple mutant genotypes

Barcodes in our libraries contain a Poisson-distributed number of mutations (Figure S1C). Though most mutations are sampled in at least one barcode as a unique single mutant (Figure S1E), most library genotypes contain multiple amino-acid mutations, and some amino-acid mutations are only sampled on many of these multiple-mutant backgrounds. Therefore, we used global epistasis models (Otwinowski et al., 2018) to decompose single mutation effects from across the set of single- and multi-mutant backgrounds (Figure S3). Briefly, we fit regression models that represent the phenotype of each library variant as a sum of latent-scale effects of all component amino-acid mutations, which are transformed by a flexible nonlinear curve to the observed experimental scale; the shape of the nonlinear curve and the single-mutant effect terms are fit simultaneously to all of the data. For variance estimates on each library variant, we used the standard error of the estimate on *K*_D,app_ to estimate a variance for our per-variant binding measurements; for expression, we calculated empirical estimates of variance as a function of cell count, based on binning replicate wildtype barcodes present in the library across bins of sampling depth. Our analysis, implemented in the `dms_variants` package (see https://jbloomlab.github.io/dms_variants/dms_variants.globalepistasis.html), is as described by Otwinowski et al., except we used a Cauchy likelihood model to relate observed measurements to the global epistasis modeled phenotype, which should be more tolerant of outliers than the Gaussian likelihood used by Otwinoski et al., and we transformed our single mutant effect latent-scale coefficients back to the experimentally measured observed scale, to facilitate comparison with additional measurements made on this scale such as the RBD homologs spiked into each library. Computational notebooks detailing the global epistasis fits are provided on GitHub (https://github.com/jbloomlab/SARS-CoV-2-RBD_DMS/blob/master/results/summary/global_epistasis_expression.md and https://github.com/jbloomlab/SARS-CoV-2-RBD_DMS/blob/master/results/summary/global_epistasis_binding.md).

For our binding titration measurements, directly measured single mutant phenotypes correlated extremely well between replicates (R^2^=0.97, Figure S3E), and this correlation was not further improved by the global epistasis decomposition (Figure S3F); therefore, we retained all directly measured single-mutant effects, and only used global epistasis decomposition to interpolate the 14% of single mutants in each library that were not directly measured on any single-mutant backgrounds (which together comprise the measurements correlated in Figure 2F). It is important to note that the shape of global epistasis nonlinearity that was fit to the data disallows mutations from increasing affinity relative to wildtype (Figures S3D, G)—this prevents us from ascribing affinity-enhancing effects to any of the mutations that we did not directly measure as single mutants (only 5.7% of mutants were not sampled as single mutants in either library), which we accept as an appropriately conservative approach.

In the case of our expression measurements, directly sampled single mutants correlated moderately well between replicates (R^2^=88, Figure S3B), but this correlation was improved between the global epistasis estimates derived from each library (R^2^ = 0.93, Figure 2E). This may be in part because the expression phenotype is a more widely distributed phenotype with smaller relative shifts in the mean caused by mutation, and because of the errant outliers that we could not account for as discussed above with regards to wildtype barcodes, such that measurements of mutational effects are improved when integrating across many different backgrounds instead of taking a single observed barcode at face value. Therefore, for expression phenotypes, we used the global epistasis estimates for all mutations. We filtered out four coefficients from library 1 and three from library 2 that had nonsensically high model estimates, likely to do partial collinearities among some low-coverage mutations. Our final binding and expression single-mutant phenotypes were determined from the average effect across the two independent library replicates. A computational notebook detailing the full derivation of our final single mutant phenotypic scores for binding and expression is on GitHub (https://github.com/jbloomlab/SARS-CoV-2-RBD_DMS/blob/master/results/summary/single_mut_effects.md#assessing-global-epistasis-models-for-binding-data).

### Data visualization

The interactive heatmap of mutational effects shown at https://jbloomlab.github.io/SARS-CoV-2-RBD_DMS/ was made using the `altair` (VanderPlas et al., 2018) Python package.

For the logo plot representation of the data in Figure S4, the experimental measurements of Δlog(MFI) and Δlog_10_(*K*_D,app_) were converted to letter heights as follows. For binding, we first computed a Boltzmann-like weighting factor for each amino acid *a* at site *r* as w_r,a_= exp(**α** x_r,a_) where x_r,a_ is the experimental measurement for the effect of the mutation of site *r* to amino acid *a*, in other words the Δlog(MFI) or Δlog_10_(*K*_D,app_) value. The **α** parameter is a temperature-like scaling factor which was set to 1.4 for the binding values, and chosen for the expression values so that the range of exponents for expression is the same as for binding. The letter heights were then computed by re-scaling the weighting factors at each site to sum to one, so that the letter height is p_r,a_ = w_r,a_ / ∑_a’_ w_r,a’_. The logo plots themselves were rendered using `Logomaker` (Tareen and Kinney, 2020). The code that creates these logo plots is on GitHub at https://github.com/jbloomlab/SARS-CoV-2-RBD_DMS/blob/master/results/summary/logoplots_of_muteffects.md.

The interactive structure-based visualizations at https://jbloomlab.github.io/SARS-CoV-2-RBD_DMS/structures were built using `dms-view` (Hilton et al., 2020). In these visualizations, the logo plot letter heights were computed as for Figure S4 (see paragraph immediately above). Number of effective amino acids was calculated as the exponentiated preferences. Mean, minimum, and maximum mutational effects per site were calculated from the set of Δlog(MFI) or Δlog_10_(*K*_D,app_) measurements of all missense mutations at a site.

### Structural analyses

Structural analyses of the ACE2-bound SARS-CoV-2 and SARS-CoV-1 RBDs used the crystal structures from PDB 6M0J (Lan et al., 2020) and 2AJF (Li et al., 2005a), respectively. ACE2 contacts were annotated as residues with any non-hydrogen atom within 4 Angstrom from any ACE2 residue. Solvent accessible surface area was calculated from the 6M0J structure using `dssp` (W Kabsch, 1983), with and without the ACE2 ligand present. Relative solvent accessibilities were determined by normalizing to the maximum theoretical solvent accessibility of a residue (Tien et al., 2013). Structural images were rendered in PyMol. Full analyses of our mutational measurements in context of structural and evolutionary features are provided on GitHub (https://github.com/jbloomlab/SARS-CoV-2-RBD_DMS/blob/master/results/summary/structure_function.md and https://github.com/jbloomlab/SARS-CoV-2-RBD_DMS/blob/master/results/summary/sarbecovirus_diversity.md).

Antibody epitopes were mapped from crystal structures 6W41 (Yuan et al., 2020b), 6WAQ (Wrapp et al., 2020b), 2DD8 (Prabakaran et al., 2006), 3BGF (Pak et al., 2009), 2GHW (Hwang et al., 2006), 7BZ5 (Wu et al., 2020), and cryo-EM structures 6NB6 and 6NB7 (Walls et al., 2019), and 6WPS (Pinto et al., 2020). RBD residues were annotated as being in an antibody epitope if any non-hydrogen atom was within 4 Angstroms of an antibody residue, with the exception of the backbone-only models of 6NB6 and 6NB7, where epitopes were defined as RBD residues with C_α_ within 8 Angstroms of any antibody residue. Our full analysis of mutational constraint in antibody epitopes is provided on GitHub (https://github.com/jbloomlab/SARS-CoV-2-RBD_DMS/blob/master/results/summary/antibody_epitopes.md).

### Analysis of circulating variants

All 31,570 spike sequences on GISAID as of 27 May 2020 were downloaded and aligned via `mafft` (Katoh and Standley, 2013). Sequences from non-human origins and sequences containing any gap characters were removed. All amino-acid mutations among GISAID sequences were enumerated. Some low-coverage spike sequences contain undetermined ‘X’ characters. We excluded any mutation from our curated set of GISAID mutations if it was solely observed on sequence backgrounds containing at least one undetermined X character in the RBD sequence; however, sequences with X characters were allowed to contribute to observations of mutation count for mutations that were observed on at least one other high-coverage RBD sequence. To characterize patterns of selection on amino-acid mutations observed among GISAID sequences, we conducted permutation tests as described in the Figure S8 legend. Our full analysis of mutational effects of circulating variants is provided on GitHub (https://github.com/jbloomlab/SARS-CoV-2-RBD_DMS/blob/master/results/summary/circulating_variants.md). We acknowledge all GISAID contributors for their sharing of sequencing data (https://github.com/jbloomlab/SARS-CoV-2-RBD_DMS/blob/master/data/alignments/Spike_GISAID/gisaid_hcov-19_acknowledgement_table.xls).

### Alignment and phylogeny

We used the curated RBD sequence set from Letko et al. (Letko et al., 2020), adding newly described RBD sequences from sarbecovirus strains RaTG13 (Zhou et al., 2020b), RmYN02 (Zhou et al., 2020a), GD-Pangolin and GX-Pangolin (Lam et al., 2020), and the additional non-Asian bat sarbecovirus isolate BtKY72 (Tong et al., 2009). RBD nucleotide sequences were aligned via `mafft` with a gap opening penalty of 4.5, and the maximum likelihood phylogeny was inferred in RAxML (Stamatakis, 2014) under the GTR model with 4 gamma-distributed discrete categories of among-site rate variation.

### Pseudotyped lentiviral particle infection assays

We selected seven single mutations from our deep mutational scanning measurements for validation of phenotypic effects in a spike-pseudotyped lentivirus assay (Crawford et al., 2020). Mutations were selected that exhibited deleterious effects on RBD expression (C432D) or ACE2 binding (L455Y, N501D and G502), no strong phenotypic effect on either binding or expression (N439K), and affinity-enhancing effects (Q498Y and N501F). These point mutations were introduced via site-directed mutagenesis (New England Biolabs E0554S) into the HDM vector containing codon-optimized SARS-CoV-2 Spike from Wuhan-Hu-1, with an upstream Kozak sequence. The full sequence of this plasmid is available at https://github.com/jbloomlab/SARS-CoV-2-RBD_DMS/blob/master/data/plasmid_maps/2736_HDM_IDTSpike_EcoKozak.gb.

Pseudotyped lentiviral particles were generated as previously described (Crawford et al., 2020). Briefly, 2.5e5 293T cells per well were seeded in 12-well plates in 1 mL D10 growth media (DMEM with 10% heat-inactivated FBS, 2 mM l-glutamine, 100 U/mL penicillin, and 100 μg/mL streptomycin). 24h later, cells were transfected using BioT transfection reagent (Bioland Scientific, Paramount, CA, USA) with 0.5 μg of the ZsGreen lentiviral backbone pHAGE2-CMV-ZsGreen-W (BEI Resources NR-52520), 0.11 μg each of the lentiviral helper plasmids HDM-Hgpm2 (BEI Resources NR-52517), pRC-CMV-Rev1b (BEI Resources NR-52519), and HDM-tat1b (BEI Resources NR-52518), and 0.17 μg wildtype or mutant SARS-CoV-2 Spike plasmids. Media was changed to fresh D10 at 24 h post-transfection. At 60 hours post transfection, the viral supernatant was collected, filtered through a 0.45 μm SFCA low protein-binding filter, and stored at -80°C. Viruses were rescued in triplicate.

The resulting viruses were titered as previously described (Crawford et al., 2020). 293T cells stably expressing ACE2 (BEI NR-52511) were seeded at 1e4 cells per well in poly-L-lysine coated 96-well plates (Greiner 655930). 24 h later, 3 wells were counted and averaged to determine the number of cells per well at time of infection. Media was removed from the 293T-ACE2 cells and replaced with fresh D10 containing 50 μL of pseudovirus supernatant in a final volume of 150 μL. Polybrene (TR-1003-G, Sigma Aldrich, St. Louis, MO, USA) was added to a final concentration of 5 μg/mL. 60 h post-infection, cells were analyzed by flow cytometry. Titers were calculated using the Poisson formula. If *P* is the percentage of cells that are ZsGreen positive, as determined by drawing a ZsGreen+ gate from uninfected controls, then the titer per ml is: *-ln(1* − *P/100) × (number of cells/well)/(volume of virus per well in mL)*. Titers are only accurate when the percentage of ZsGreen+ cells is relatively low, i.e., ∼1-10%. Titers are reported relative to the mean of the wildtype, which had similar titers as Crawford et al. of ∼10^4^ infectious particles per mL (Crawford et al., 2020). The dashed horizontal line in Figure 4E showing the limit of detection was calculated as the minimum titer that would be determined in the case of a single positive event.

### RBD purification and binding assays

The RBDs of SARS-CoV-1, SARS-CoV-2, WIV1, RaTG13, SHC014, ZC45, and ZXC21 were synthesized by GenScript and cloned into vector pCMV with a preceding mu-phosphatase signal peptide and a terminal octa-histidine tag. Plasmids were transfected into 150mL suspension expi293F or HEK293F cells at 37°C in a humidified 8% CO2 incubator rotating at 130 rpm and harvested 3 days later. Clarified supernatants were purified in batch over Talon resin (Takara) prior to buffer exchanging into 20mM Tris pH8 150mM NaCl and flash freezing.

Biolayer interferometry binding assays were performed on an Octet Red instrument at 30°C with shaking at 1,000 RPM. ARG2 biosensors were hydrated in water then activated for 300 s with an NHS-EDC solution (ForteBio) prior to amine coupling. 5-10 µg/mL of each RBD was loaded in a buffer containing 10mM pH5 sodium acetate onto ARG2 tips (ForteBio) for 600 seconds and then quenched into 1M ethanolamine for 600 seconds. A baseline in 10X kinetics buffer (ForteBio) was collected for 120 s prior to immersing the sensors in a 1:3 serial dilution of his-tagged human ACE2 (Sino Biologicals) ranging from 1,000 to 0.83nm in 10X Kinetics Buffer. Curve fitting was performed using a 1:1 binding model and the ForteBio data analysis software. Mean *k*_on_, and *k*_off_ values were determined with a global fit applied to all data.

---

**Figure S1.**
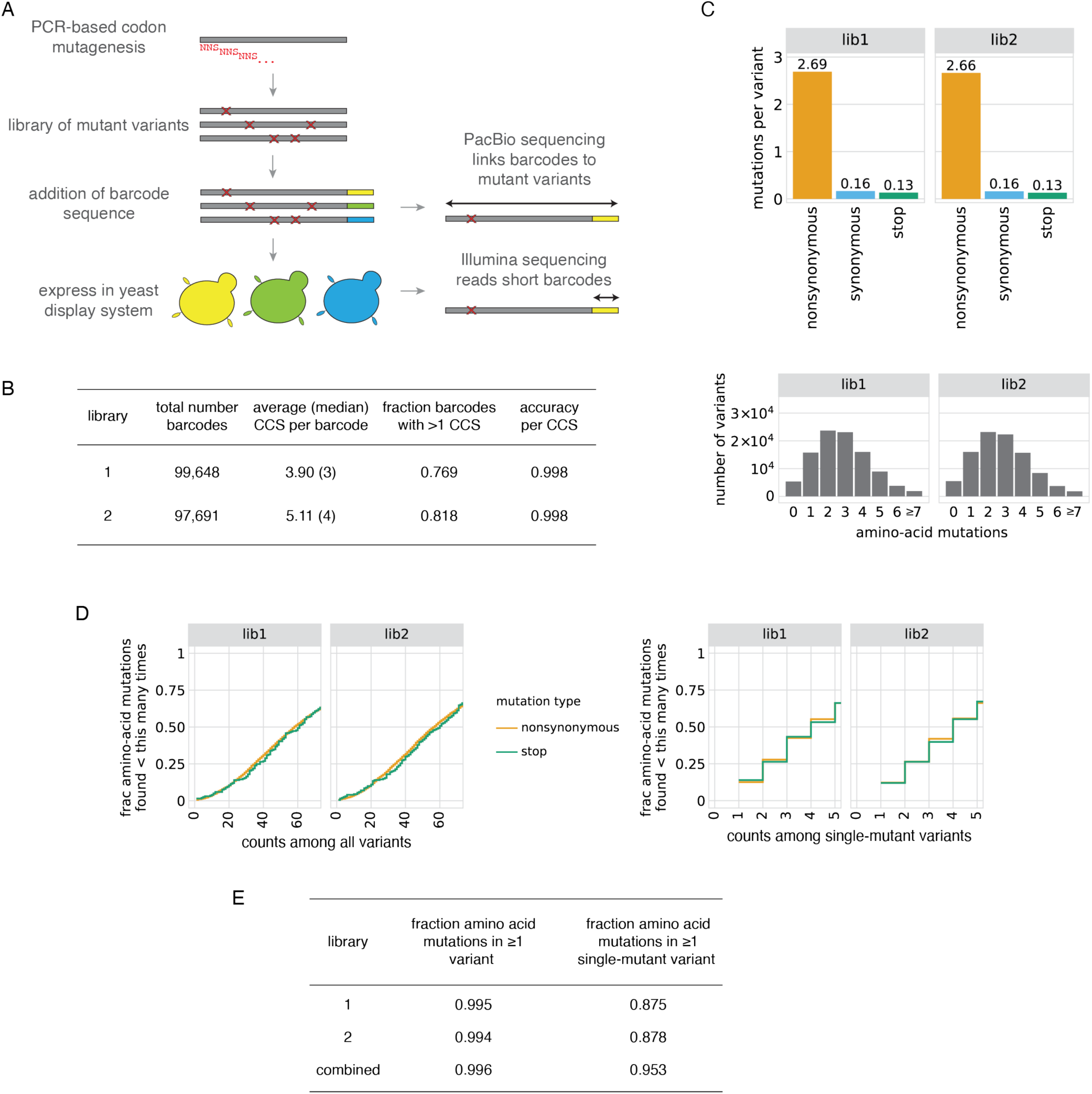
SARS-CoV-2 RBD mutant libraries. (A) Scheme of the library generation and sequencing approach. SARS-CoV-2 RBD mutant libraries were constructed in fully independent duplicates, and variants were linked to barcodes by long-read PacBio sequencing. (B) PacBio sequencing stats on duplicate SARS-CoV-2 mutant libraries. Comparison of RBD sequences among independent circular consensus sequences (CCSs) of the same barcode enables calculation of an empirical accuracy, which describes the minimal expected accuracy of the barcode:RBD linkage for barcodes with a single CCS (see Methods for details). Most barcodes were represented by multiple CCSs, which further increases the accuracy of barcode:RBD linkage. (C) Statistics on mutation rates in mutant libraries. Top, average number of mutations of different types across variants in each library. Bottom, distribution of number of amino-acid mutations per variant. (D, E) Mutation coverage in mutant libraries. Cumulative distribution plots (D) give the fraction of all possible amino-acid mutations observed in the indicated number of variants, including all variants (left) or only variants with a single mutation (right). Minimum coverage statistics from these curves are tabulated in (E).

**Figure S2.**
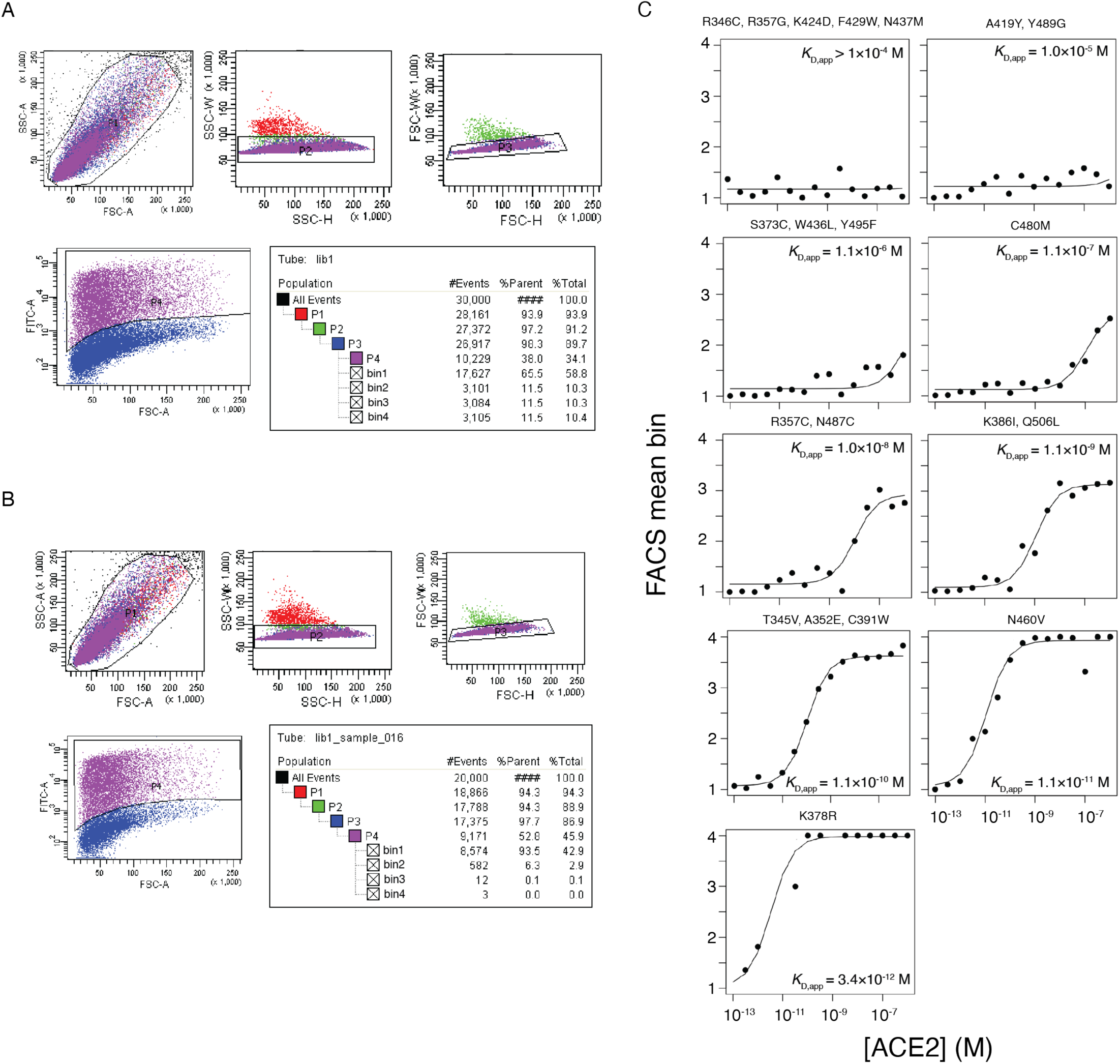
FACS-based determination of variant phenotypes. (A, B) Representative sorting gates used to select cells for for expression (A) and binding (B) FACS experiments. FSC and SSC gates select for single cells (P1-P3), and FITC labeling of an RBD C-terminal epitope tag defines RBD+ gates (P4), when necessary. Tables show the nested hierarchy of sort gates, with final bins 1-4 for expression and binding shown in Figure 2A and 2B, respectively. For (A), the P4 “RBD+” gate was used to enrich the library for expressing variants, which were grown up and re-induced for binding experiments as in (B). (C) Example variant-specific titration curves inferred from the deep mutational scanning experiment. Randomly sampled titration curves are illustrated across the range of fit *K*_D,app_ binding constants, with variant genotype listed above each panel. Because curves that were fit with *K*_D,app_ between 10^−4^ to 10^−6^ were virtually indistinguishable non-responsive curves, we truncated all *K*_D,app_ measurements in this range to a censored >10^−6^ M cutoff.

**Figure S3.**
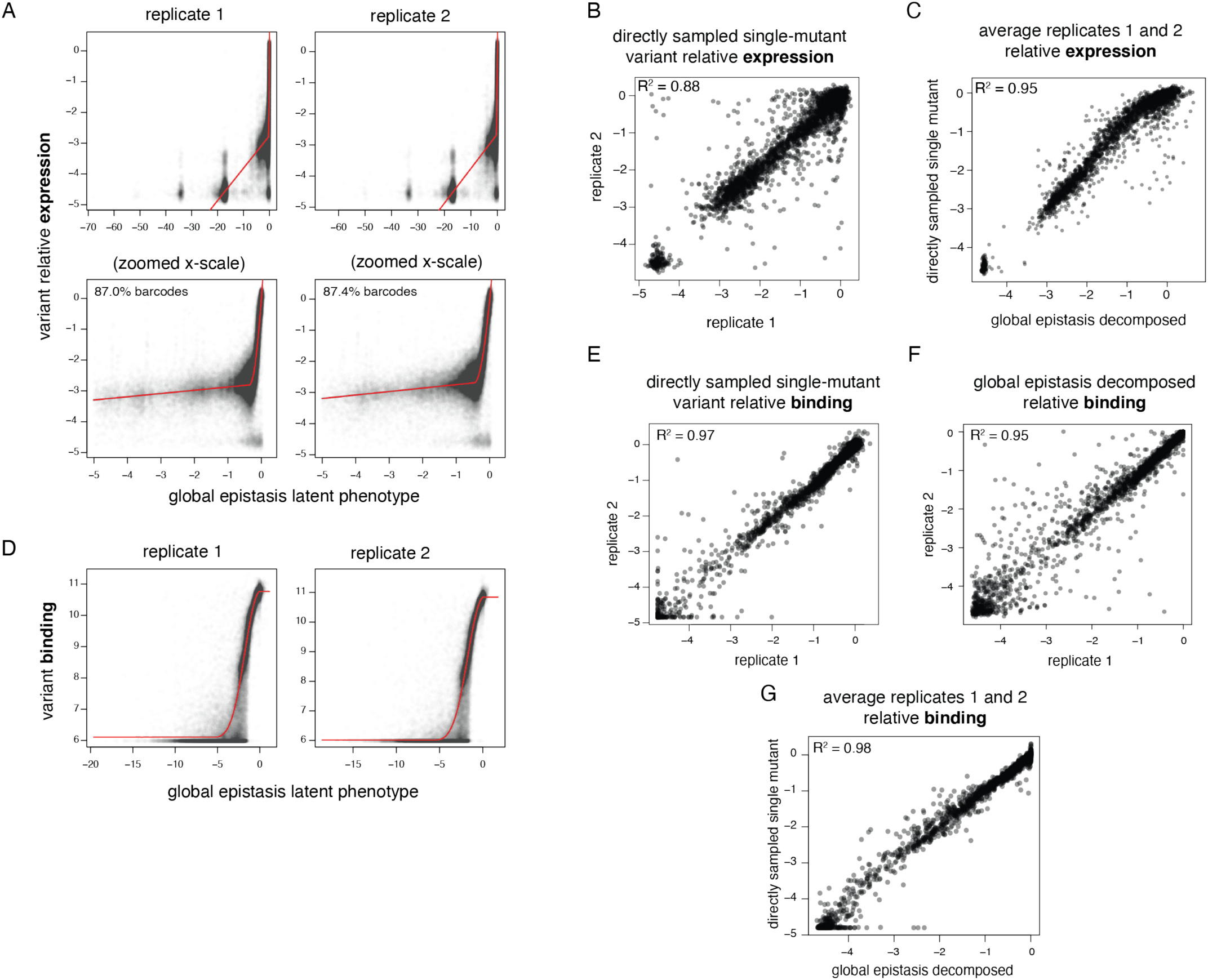
Global epistasis decomposition of single-mutant effects. Global epistasis models were fit to decompose single-mutant effects from variant backgrounds containing variable numbers of mutations. These models invoke an underlying latent scale on which mutations combine additively, which is linked to the experimental scale by a flexible nonlinear curve fit, which accounts for limits in dynamic range and other nonlinearities. See the Methods for more details. (A, D) Global epistasis fits. Plots illustrate, for each library variant, its experimentally determined phenotype for expression (A) or binding (D) versus its latent phenotype predicted by the global epistasis model. Red lines indicate the shape of the nonlinear curve fit. (B, E) Correlation in mutation effects on expression (B) and binding (E) between replicates, for mutations that were sampled directly as single mutants with no global epistasis decomposition. (F) Correlation in mutation effects on binding between replicates, for all global-epistasis-decomposed single-mutant effect terms on the observed phenotype scale. Equivalent plot for expression is Figure 2E. (C, G) Correlation in mutation effects on expression (C) and binding (G) averaged across replicates, for directly sampled single-mutant measurements versus global-epistasis-decomposed mutation effects. For expression, global epistasis averaging of single-mutant effects across all variants (Figure 2E) improved replicate correlations beyond the directly sampled measurements (B), so global-epistasis-decomposed values were used for all single-mutant terms. For binding, directly sampled single-mutant effects (E) were better correlated than the values decomposed from global epistasis models (F), so global epistasis models were used to interpolate single-mutant measurements only for mutations that were not observed on any directly-sampled single-mutant variant backgrounds.

**Figure S4.**
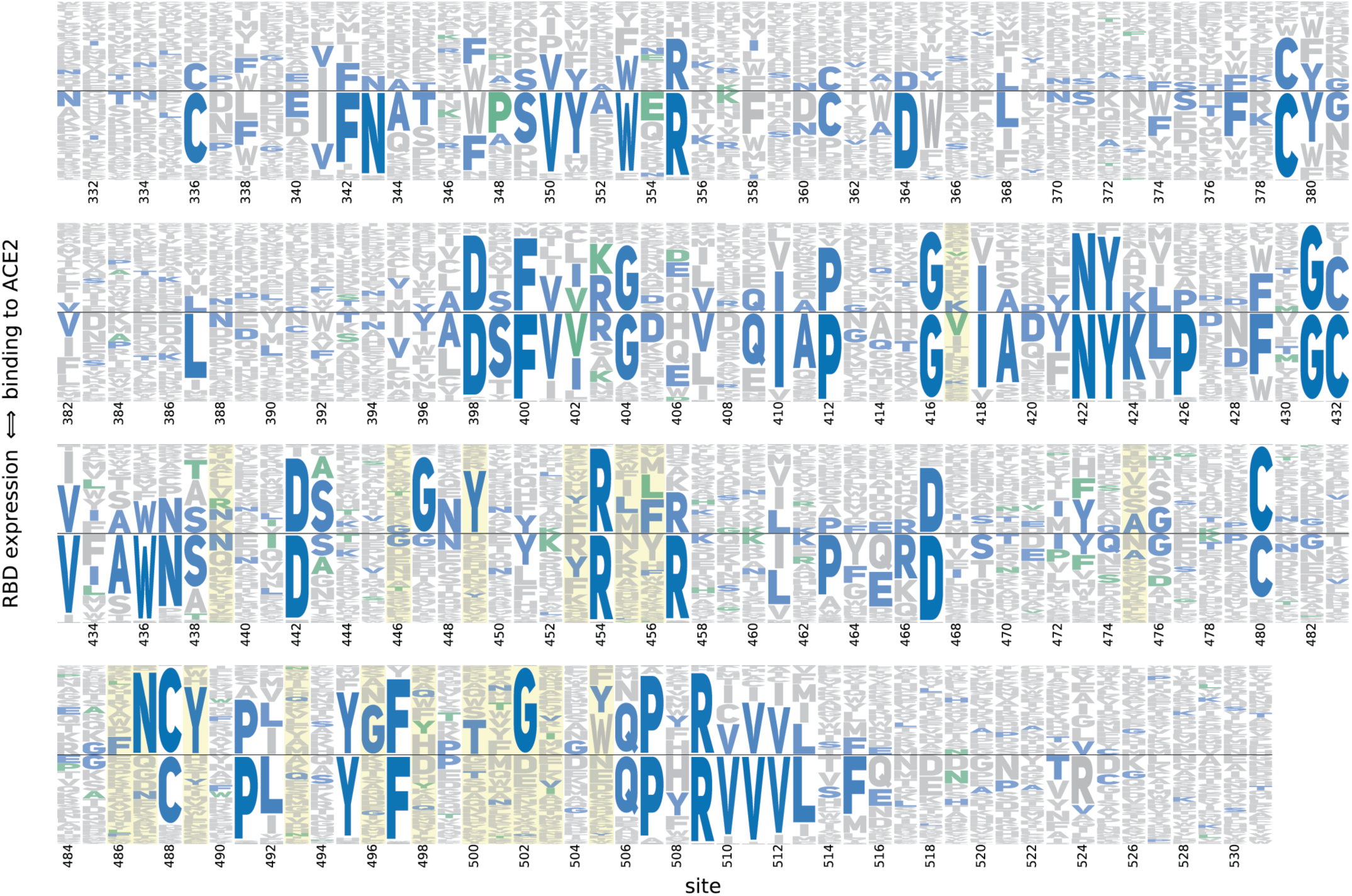
Logo plot representation of mutational effects on binding and expression. Letter height indicates preference of each site for individual amino acids with respect to ACE2 binding (height above the center line) or RBD expression (height below the center line). Blue letters indicate the unmutated SARS-CoV-2 amino acid, and, where applicable, green letters indicate differences found in SARS-CoV-1. Yellow highlights mark residues that contact ACE2 in the SARS-CoV-2 or SARS-CoV-1 crystal structures. See the Methods for details of how the amino-acid preferences are calculated from the experimental measurements.

**Figure S5.**
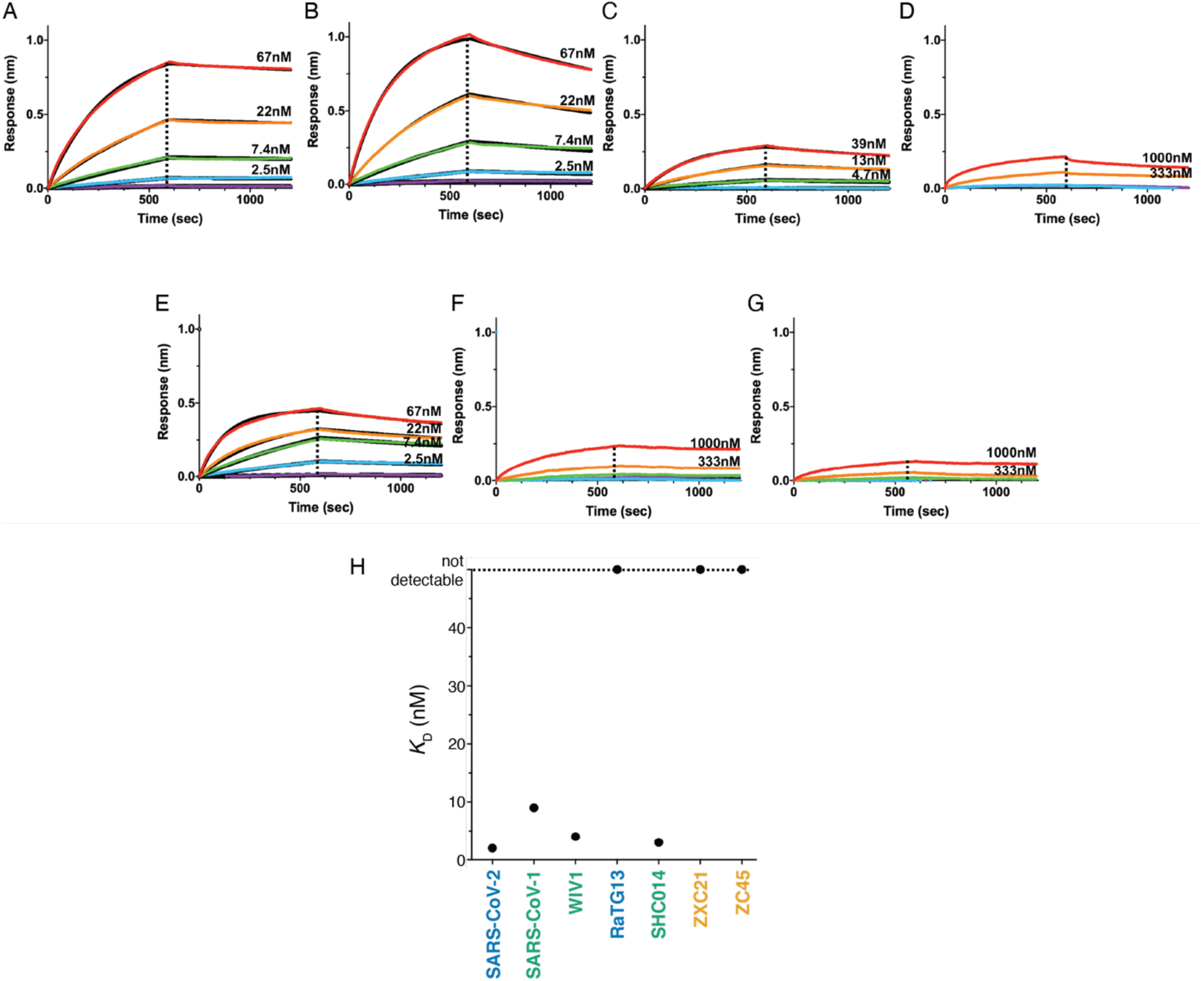
Human ACE2 binds to various sarbecovirus RBDs with distinct affinities. (A-G) Biolayer interferometry binding of various concentrations of monomeric ACE2 to the RBD of SARS-CoV-2 (A), SARS-CoV-1 (B), WIV1 (C), RaTG13 (D), SHC014 (E), ZXC21 (F) and ZC45 (G) immobilized at the surface of biosensors. Global fit curves are shown as black lines. The vertical dashed lines indicate the transition between association and dissociation phases. (H) Summary of *K*_D_ values determined from the shown BLI traces.

**Figure S6.**
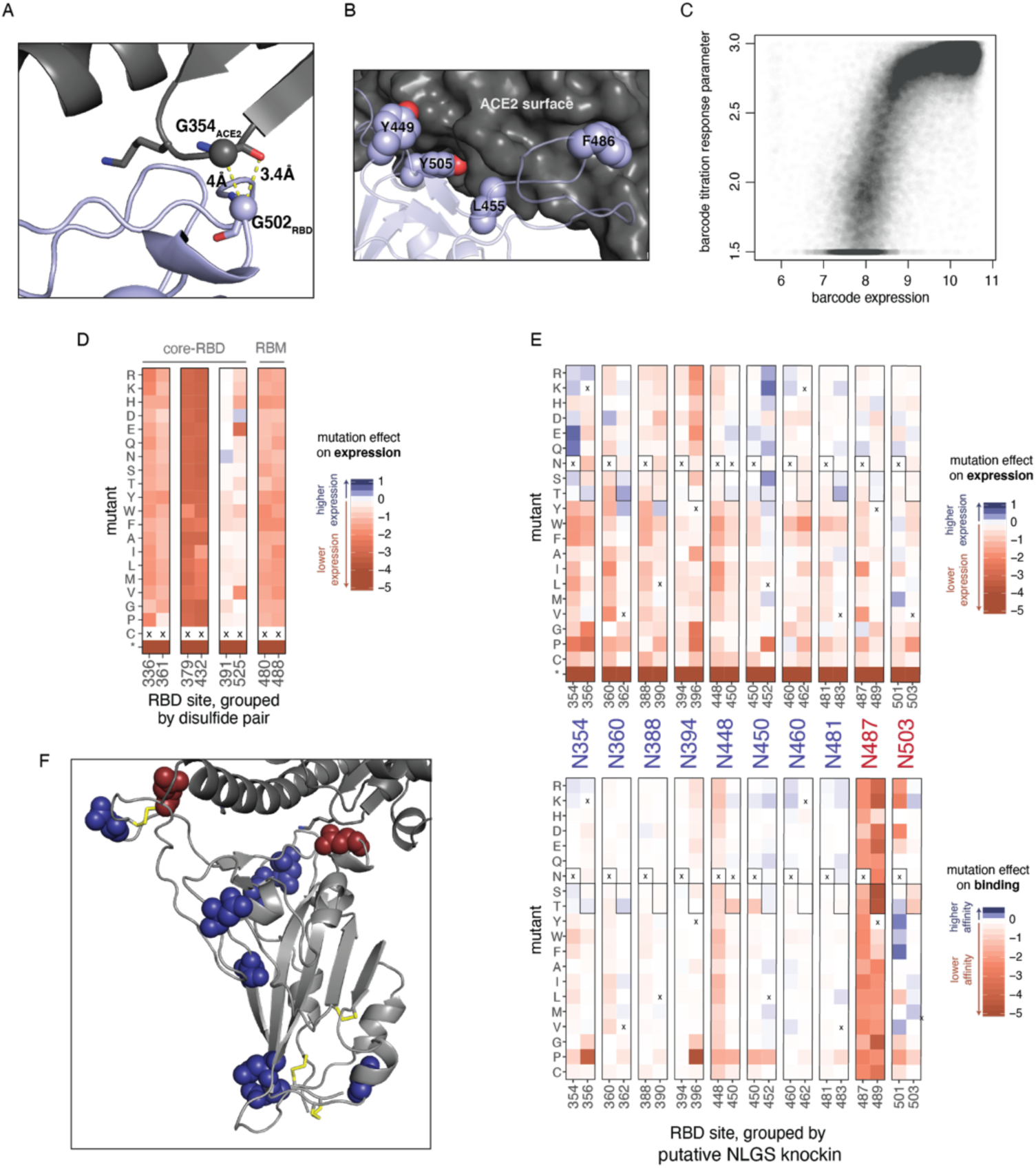
Additional structural analyses of mutation effects. (A, B) Structural depictions of sites exhibiting stability-binding tradeoffs. (A) RBD residue G502 requires small amino acid side chains for ACE2 binding (Figure 3B), consistent with its close proximity to G354_ACE2_ in the bound structure. (B) Mutations to polar residues at positions Y449, L455, F486, and Y505 would enhance expression but reduce binding, consistent with specific geometric constraints imposed by the close packing of these residues at the ACE2 surface. (C) Relationship between barcode expression and titration response plateau parameters. The correlation between mutation effects on binding and expression in Figure 5C could emerge from trivial correlation between phenotypes (e.g. yeast with higher RBD surface expression can bind more ACE2). However, our multiple-concentration titration approach should in principle remove this trivial correlation (Adams et al., 2016), because each binding phenotype is determined from a self-referenced titration curve, for which the free plateau response parameter can vary to account for different levels of saturated binding due to RBD expression (see Figure S2C). Consistent with this premise, the response parameter from the titration fit for each library variant correlates with its expression phenotype. (D) Mutation effects on expression at disulfide cysteine residues. Details as in figure 5E. (E) Effects of putative N-linked glycosylation site (NLGS) knock-in mutations. Heatmap details as in Figure 5F. There are 10 surface-exposed asparagines for which RBD expression is unaffected or enhanced (top) when an NLGS motif is introduced via mutations to S or T at the i+2 site; for eight of these putative NLGS knock-ins (blue labels), the putative glycan is also tolerated for ACE2 binding (bottom), but for two (red labels), introduction of the NLGS motif is not tolerated for ACE2 binding. (F) Mapping of these ten asparagines to the RBD structure illustrates that these two binding-constrained asparagines (red) cluster to the ACE2 interface.

**Figure S7.**
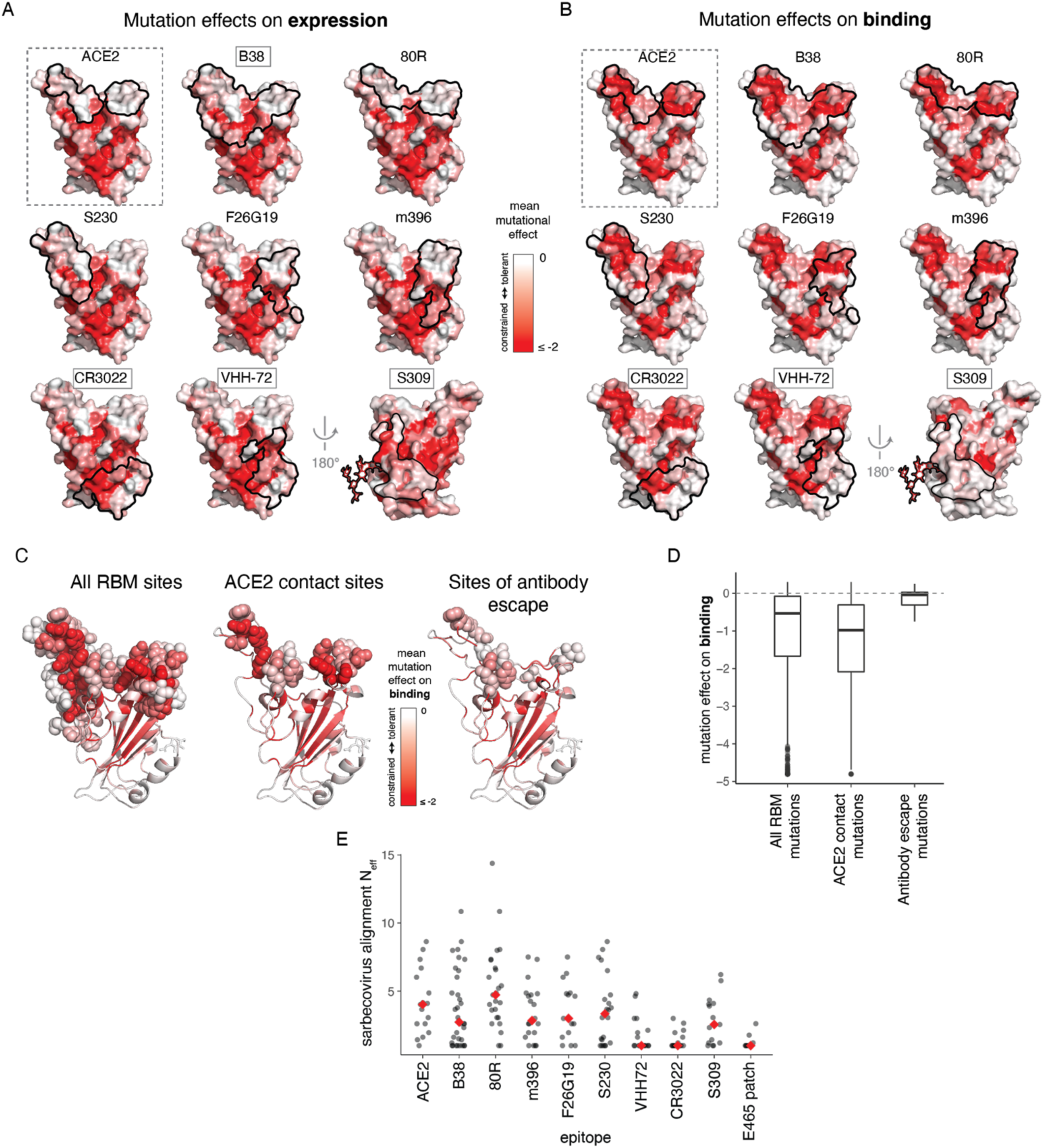
Mutational and evolutionary constraint of antibody epitopes. (A, B) Surface representations of antibody epitopes colored by mutational effects on expression (A) and binding (B). Representations as described in Figure 7A. (C,D) Mutational constraint and observed antibody escape mutations. Baum et al. (Baum et al., 2020) selected SARS-CoV-2 escape mutations from RBD-directed antibodies. We compare the average mutational tolerance of the sites at which these escape mutations accrue (C), and the effects of the specific escape mutations themselves (D) to all RBM and ACE2-contact sites/mutations. The antibody escape involved mutations that were better tolerated than typical mutations in the RBM or ACE2-binding interface. (E) Evolutionary diversity in antibody epitopes and our newly described E465-centered surface patch among the sarbecoviruses in Figure 1A. Diversity is summarized as the effective number of amino acids (N_eff_), which scales from 1 for a site that is invariant, to 20 for a site in which all amino acids are at equal frequency.

**Figure S8.**
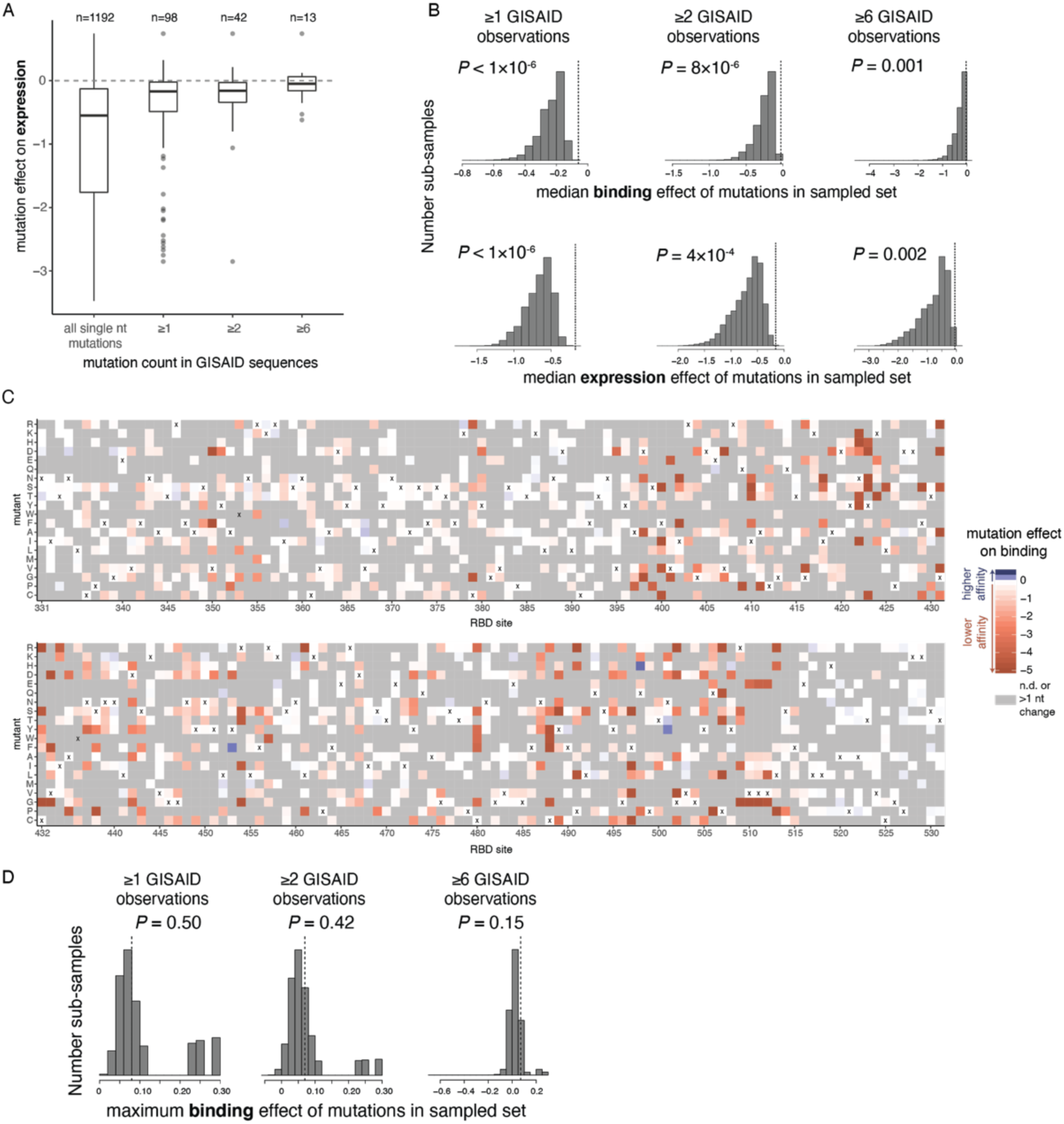
Genetic variation and selection in SARS-CoV-2. (A) Distribution of expression effects of mutations observed among circulating SARS-CoV-2 isolates. Details as in Figure 8A. (B) Permutation tests indicating the action of purifying selection on binding (top) and expression (bottom) among circulating SARS-CoV-2 mutations. For each threshold of GISAID observation counts, 1 million random sub-samples of single-nucleotide-accessible amino acid changes were generated at the same sample size as the true mutation set (n=98, 42, and 13 for the ≥1, ≥2, and ≥6 thresholds). A *P*-value was determined as the fraction of sub-samples with median mutational effect on binding or expression equal to or greater than that of the actual GISAID mutation set (dashed vertical line). The observation that the set of mutations observed in GISAID have a more favorable median mutational effect on binding and expression than randomly sampled mutations indicates the action of purifying selection for ACE2 binding and RBD stability. (C) Heatmaps depicting effects of mutations on ACE2 binding, indicating only those mutations that are accessible via single-nucleotide mutation from the SARS-CoV-2 Wuhan-Hu-1 isolate gene sequence. Amino-acid mutations that require more than one nucleotide change are in gray. (D) Permutation tests for positive selection for enhanced ACE2 affinity. Random sub-samples were generated as in (B), and the maximum affinity-enhancing effect of mutations in each sub-sample was compared to that in the actual GISAID mutation set. A *P*-value was determined as the fraction of sub-samples with a maximum effect on binding equal to or greater than in the actual GISAID mutation set (vertical dashed line). We do not see evidence for selection for enhanced ACE2 binding, as randomly sampled mutations generally contain mutations with stronger affinity-enhancing effects than observed in the GISAID mutation set.

## List of Supplemental Files

- Supplemental File 1: CSV containing binding and expression measurements for RBD homologs spiked into the deep mutational scanning libraries
- Supplemental File 2: html file of the interactive heatmap provided via url in the main text
- Supplemental File 3: CSV file containing all single-mutant deep mutational scanning measurements from our duplicate experiments

