## Supplemental File 2 for "Deep mutational scanning of SARS-CoV-2 receptor binding domain reveals constraints on folding and ACE2 binding"

Instructions | SARS-CoV-2 RBD DMS


### SARS-CoV-2 RBD DMS

##### Instructions

- Hover over cells with mouse to reveal additional information.
- Select site subsets using the drop down menu below the plots.
- Change which sites are displayed by brushing the zoom bar and dragging the brush.
- Clear the zoom bar by double clicking it.
- Structural visualizations of the data are available via `dms-view` here
- Raw data available on GitHub
